# Tiarins, a diverse family of natural Trojan-horse aminoacyl-tRNA synthetase inhibitors discovered by genome mining

**DOI:** 10.64898/2025.12.02.691738

**Authors:** Dmitrii Y. Travin, Aleksei Livensky, Dmitry Bikmetov, Constantine Pavlov, Marina Serebryakova, Konstantin Gilep, Fengjie Wu, Delphine Naquin, Tatiana Timchenko, Guy Lippens, Séverine Zirah, Peter Mergaert, Svetlana Dubiley

## Abstract

Trifolitoxin (TFX) is a ribosomally synthesized and post-translationally modified peptide antibiotic produced by *Rhizobium anhuiense* T24. Although discovered more than half a century ago, its mechanism of action has remained elusive. Here we demonstrate that TFX inhibits arginyl-tRNA synthetase (ArgRS), an essential translation enzyme. TFX acts as a Trojan-horse antibiotic: it enters cells via the inner membrane oligopeptide transporter YejABEF and then undergoes partial proteolysis by the aminopeptidase PepN to release a “warhead” that binds ArgRS and arrests translation. A systematic analysis of prokaryotic genomes revealed that TFX belongs to a widespread family of modified peptides that we designate the tiarins. In addition to ArgRS inhibitors, the tiarin family includes compounds that specifically target at least seven other aminoacyl-tRNA synthetases. These findings define a previously unrecognized family of natural antibiotics and provide a framework for developing a new class of inhibitors targeting various tRNA synthetases.

## INTRODUCTION

Ribosomally synthesized and post-translationally modified peptides (RiPPs) comprise a rapidly expanding class of natural products. They share a common biosynthetic logic, in which a precursor peptide undergoes biochemical transformation by specific enzymes, installing posttranslational modifications (PTMs)^1^. Typically, the precursor contains an N-terminal leader peptide part that serves as a docking site for the RiPP Recognition Element (RRE) domain of PTM-installing enzymes and a core part that is converted into the mature compound. The genes encoding the precursor(s), PTM-installing enzymes, and other associated proteins, such as export pumps, transcription factors, and self-immunity proteins, often form biosynthetic gene clusters (BGCs) in bacterial genomes, facilitating their coordinated regulation and horizontal transfer. The biological roles of RiPPs are exceptionally diverse, encompassing cofactors, metallophores, signal molecules, antivirals, antimicrobials targeting various cellular processes, *etc*.^2,3^. Although advances in genome mining techniques have led to an explosion in the number of newly characterized RiPP families, only a small subset has experimentally validated bioactivities and identified molecular targets. Furthermore, the detailed mechanisms of action have been elucidated for an even smaller fraction of these compounds.

Trifolitoxin (TFX), a RiPP antibiotic produced by *Rhizobium anhuiense* bv. *trifolii* T24 (previously classified as *R. trifolii* and *R. leguminosarum* bv. *trifolii*)^4,5^, inhibits the growth of various Alphaproteobacteria, including symbiotic (*Rhizobium, e.g.*, *R. leguminosarum* bv. *phaseoli* 4292 (**Fig. 1a**) and free-living diazotrophs (e.g., *Rhodospirillum*), as well as plant (*Agrobacterium*) and animal (*Brucella*) pathogens^6^. TFX is an 11-amino acid-long peptide with the sequence DIGGSRQGCVA, in which internal residues are post-translationally modified^7^. According to the structure proposed by Lethbridge *et al*.^8^, Cys9 is converted into a thiazoline, while Arg6, Gln7, and Gly8 are modified to form a UV-absorbing fluorophore (**Fig. 1b**) containing a 1,2,4,5-substituted imidazole ring. The acylimine bond between Ser5 and Arg6 is spontaneously hydrated to give the hydroxamic acid^8^ (**Fig. 1c,d**). Despite more than five decades of study^4^, the mechanism underlying TFX bioactivity has remained unknown.

**Figure 1.**
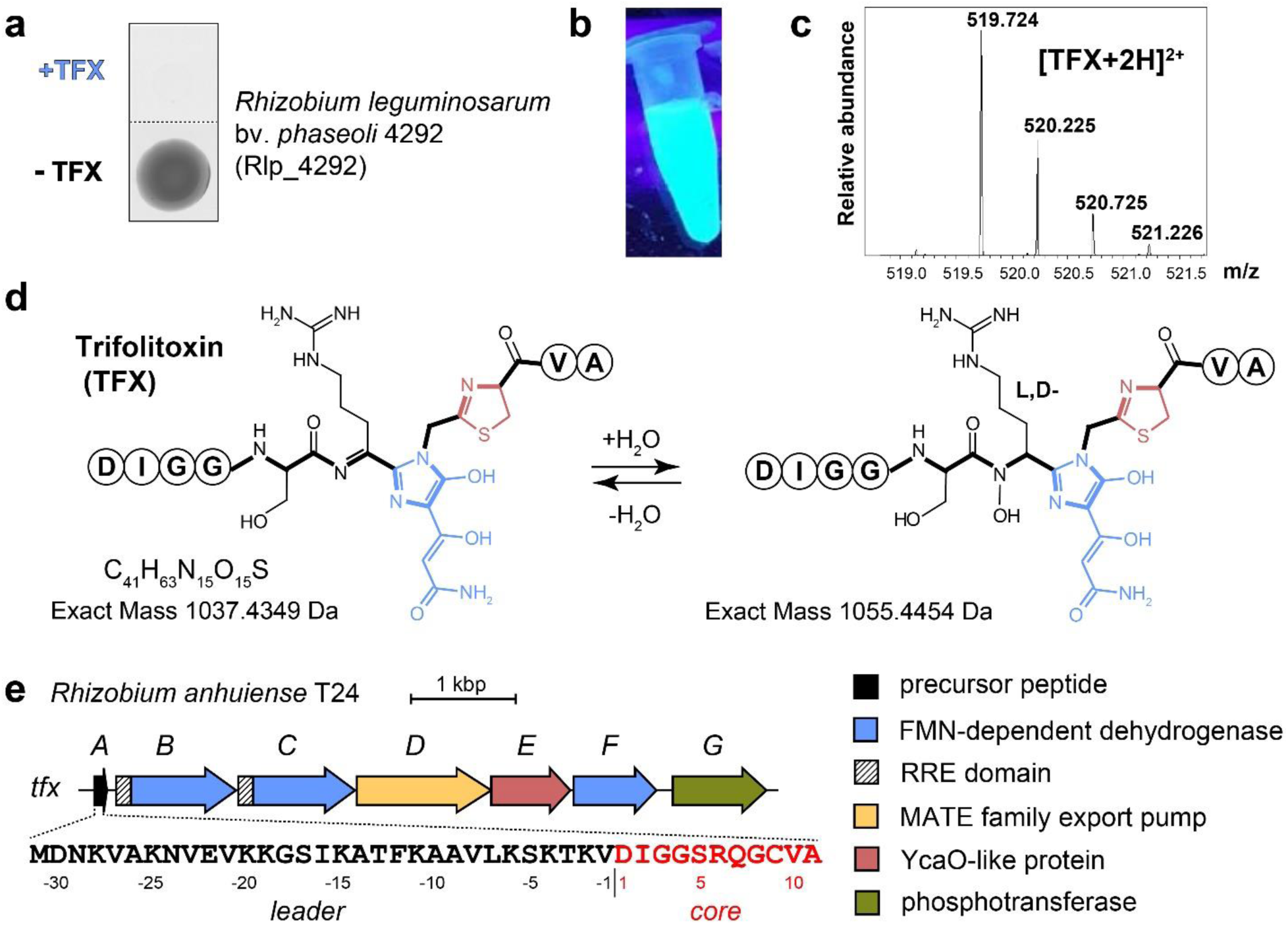
Trifolitoxin and the *tfx* biosynthetic gene cluster. **a.** Growth of TFX-susceptible strain *Rhizobium leguminosarum* bv*. phaseoli* 4292 on a medium without TFX and with TFX added. **b**. Blue fluorescence of purified TFX under the UV light (360 nm). **c.** High resolution ESI-MS spectrum of mature TFX. The *m/z* values for each peak are indicated. Theoretical MW: 1037.4349; experimental MW: 1037.4335; mass error (ppm): 1.35. **d.** Structure of mature TFX according to Lethbridge *et al*.^8^. Thiazoline cycle is shown in pink, the UV-absorbing chromophore in blue. The peptide backbone is shown in bold. **e.** Schematic representation of the *tfx* biosynthetic gene cluster from the genome of *Rhizobium anhuiense* T24. The amino acid sequence of the TfxA precursor peptide is shown with core and leader sequences highlighted in red and black, respectively. The functional annotation of the encoded proteins are listed on the right. FMN – flavin mononucleotide, MATE – multidrug and toxic compound extrusion, RRE – RiPP recognition element.

Here we demonstrate that TFX is a selective inhibitor of arginyl-tRNA synthetase (ArgRS). Moreover, we show that the antibiotic acts through an elegant Trojan-horse mechanism: TFX exploits the inner membrane oligopeptide transporter YejABEF to enter the bacterial cytoplasm, where a non-specific PepN aminopeptidase processes its unmodified N-terminal part. The released modified C-terminal portion of TFX acts as a toxic “warhead” that binds ArgRS, inhibiting its activity and thereby arresting protein synthesis. By genome mining, we uncovered a diverse family of trifolitoxin-like compounds that share the common mode of action but target different types of aminoacyl-tRNA synthetases (aaRSs). We designate this new family of RiPPs as tiarins <u>(*t*</u>rifolitoxin-like *i*nhibitors of *<u>a</u>*minoacyl-t*<u>R</u>*NA synthetases) and predict molecular targets for each identified representative.

## RESULTS

### Trifolitoxin is a member of a diverse family of natural products

The trifolitoxin BGC (*tfx*) (**Fig. 1e, Supplementary Fig. 1**) contains the genes of the 42-aa-long precursor peptide TfxA, the export pump TfxD, and putative modification enzymes^9^. Although the biosynthetic pathway of TFX remains unknown, it was proposed that the YcaO-like enzyme TfxE installs the thiazoline heterocycle, while the putative FMN-dependent dehydrogenases TfxB, TfxC, and TfxF contribute to the fluorophore formation^8^. A gene coding for a phosphotransferase, *tfxG*, whose role in TFX biosynthesis remains unclear, is located downstream of the *tfxABCDEF* operon^10^. No similarly organized clusters encoding the biosynthesis of putative TFX homologs have been described before, making the *tfx* BGC first of a kind.

To explore the diversity of TFX-like compounds, we searched for *tfx*-like BGCs across the bacterial and archaeal genomes. We focused on the putative FMN-dependent dehydrogenases TfxB and TfxC (IPR020051 and IPR000415), as they are proposed to catalyze the installation of the family-defining PTM, the blue fluorophore^8^. Consistent with their primary role in the TFX biosynthesis, TfxB and TfxC are the only enzymes in the *tfx* BGC that harbor RRE domains^11^ (**Fig. 1e**), suggesting that they perform the initial leader peptide-dependent steps in the TfxA posttranslational modification.

The search for TfxC homologs associated with TfxB proteins in prokaryotic genomes from the NCBI RefSeq database, followed by clustering with MMseqs2^12^ (at 95% identity cutoff), yielded 79 non-redundant representative sequences, which we used for a phylogenetic tree construction (**Fig. 2a** and **Supplementary Fig. 2**). The identified homologs were present in organisms spanning two archaeal and seven bacterial phyla, with the largest numbers of biosynthetic gene clusters originating from Actinomycetota (35 hits) and Pseudomonadota (24 hits) genomes (**Fig. 2a**, “Phylum” column). To identify core operons of the retrieved BGCs, we examined the ORFs located within 200 bp from each other and transcribed in the same direction as *tfxC* homologs.

**Figure 2.**
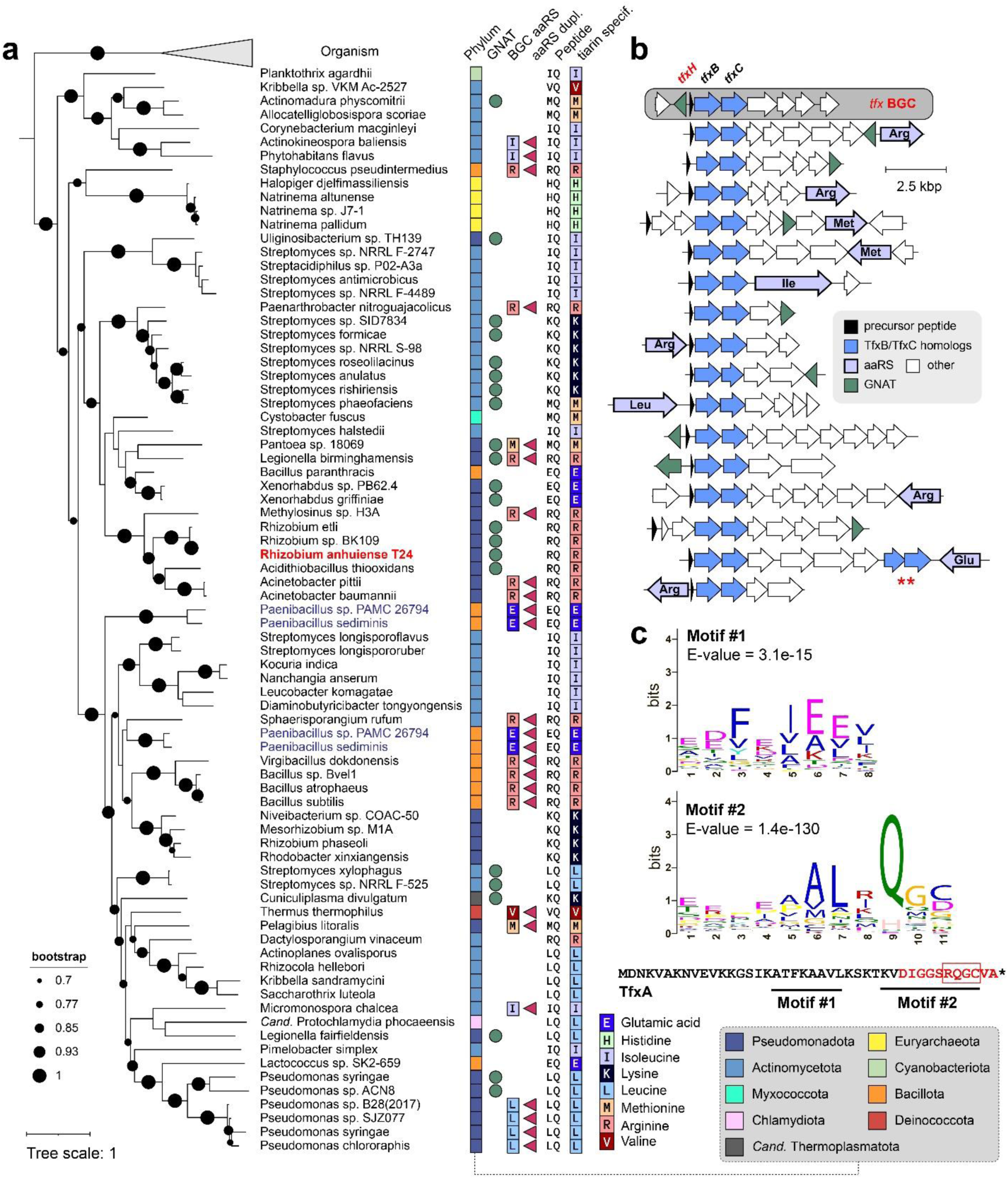
Genome mining-guided discovery of tiarins. **a.** Phylogenetic tree of TfxC homologs (n = 79). Dataset consisted of representative sequences after MMseqs2 clustering with 95% identity cutoff. For each protein sequence, the source organism and its phylum (column “Phylum”, see the color code on the right) are shown. The presence of a GNAT in the corresponding BGC is indicated with a green circle (column “GNAT”). When a BGC encodes an aaRS, its specificity is shown using a single-letter code (column “BGC aaRS”). The column “Peptide” shows the amino acid residue preceding the conserved Q in the precursor’s core sequence. A red triangle indicates the presence, elsewhere in the genome, of a duplicated copy of the aaRS encoded by the corresponding BGC (column “aaRS dupl.”). The predicted specificity of the tiarin is indicated in the column “tiarin specif.”. TfxC from *R. anhuiense* is shown in bold red. TfxC homologs from *Paenibacillus* sp. PAMC 26794 and *P. sediminus* (shown in blue) appear twice in the tree because their BGCs contain a duplicated *tfxBC* homolog pair. **b.** Representative *tfx*-like BGCs identified across bacterial and archaeal genomes, visualized with clinker^48^. Each arrow represents a gene; the proposed functions of encoded proteins are listed. The specificities of the aaRSs encoded are labeled. A double red asterisk marks a duplication of the *tfxBC*-homologs pair. **c.** Sequence logos of motifs in the predicted precursor peptides from the *tfx*-like BGCs and their locations in the amino acid sequence of the TFX precursor peptide TfxA. The leader peptide is shown in black, the core peptide is red, and residues undergoing posttranslational modification are boxed.

Next, we looked for ORFs encoding putative precursor peptides (**Supplementary Data 1)**. Most predicted precursors were encoded immediately upstream of the *tfxB-tfxC* homologs pair (**Fig. 2b**) and were associated with strong Shine-Dalgarno sequences^13^. The putative precursors varied considerably in length (23-65 amino acid residues) and, at first glance, had little to no sequence similarity. However, the search for consensus using MEME^14^, revealed two motifs having notable conservation over the whole set of predicted precursor sequences (**Fig. 2c)**. Motif #1, which shows weak sequence conservation, includes predominantly aliphatic and negatively charged amino acids. It is located near the N-terminus of the peptides and maps to the leader region of Tfx (**Fig. 2с**). We suppose that it represents the consensus within the leader peptide recognized by the RRE domains of TfxB and/or TfxC. The consensus of motif #2 includes a highly conserved Gln residue, which, in the case of TfxA, is the residue undergoing the conversion into the blue fluorophore characteristic of TFX.

Analysis of protein domains (**Supplementary Data 2**) revealed that approximately half of the putative core operons code for one to three FMN-dependent dehydrogenases in addition to the TfxB and TfxC homolog pair. Other proteins frequently encoded in these operons include putative modification enzymes, such as YcaO-like proteins, ThiF-like adenylyl transferases, methyltransferases, oxygenases, *etc*. (**Supplementary Data 2**. However, none of these additional protein domains were present in all operons, suggesting that these enzymes install accessory modifications that contribute to the considerable structural diversity of TFX-like compounds.

### Bioinformatic analysis of tfx-like BGCs suggests a mechanism of TFX action

Expression of regulatory genes, export pumps, and immunity genes in RiPP BGCs is often regulated independently, and, therefore, these genes can be located outside the core operons formed by precursor(s) and biosynthetic genes. To define the composition of putative *tfx*-like BGCs, we analyzed the functional domain architecture of proteins encoded in the vicinity of identified core operons (two ORFs upstream and two downstream of an operon) using the Conserved Domain Database (CDD)^15^. We found that genes encoding GCN5-related N-acetyltransferases (GNATs) are frequently associated with the core operons (**Fig. 2a,b, Supplementary Data 2**). Notably, a previously overlooked gene encoding a GNAT, which we further refer to as *tfxH,* is located 191 bp upstream of the *tfxA* and transcribed in the opposite direction (**Fig. 2b**). *tfxH* is dispensable for the TFX biosynthesis^9^ and is regulated independently of the *tfx* operon, suggesting that this GNAT can perform an immunity function.

We noted that genes coding for various aaRSs were remarkably enriched near *tfx*-like operons (**Fig. 2a**, column “BGC aaRS”, **Supplementary Data 2**). aaRS-encoding genes are found in BGCs of some natural products, which are either specific aaRS inhibitors themselves (*e.g.*, indolmycin^16^) or have tRNA-dependent steps in their biosynthesis (*e.g.*, valanimycin^17^). To assess the relevance of the observed enrichment of aaRSs in the *tfx*-like BGCs neighborhood, we performed an additional search for the aaRS-encoding genes in the genomes of the strains that contain the identified BGCs. In every case, a duplicate copy of an aaRS of the same amino acid specificity was present elsewhere in the genome in addition to the homolog associated with the operon (**Fig. 2a**, column “aaRS dupl.”, **Supplementary Table 1**), suggesting that functions of the BGC-associated aaRS genes are likely related to the TFX-like compounds’ biosynthesis or resistance. We further refer to the enzymes encoded outside of the identified BGCs as housekeeping aaRSs (aaRS^HSK^) to distinguish them from those encoded in the BGCs (aaRS^BGC^).

A striking pattern emerged when we compared the specificity of the aaRSs^BGC^ with the sequences of the cognate precursor peptides. In every case, when a *tfx*-like BGC encoded an aaRS, its specificity matched with the identity of the residue immediately preceding the conserved Gln in the precursor peptide (**Fig. 2a**, columns “Peptide” and “BGC aaRS”, **Supplementary Fig. 3**). Based on this observation, we conceive that: (i) TFX-like compounds, which we termed *tiarins* (*t*rifolitoxin-like *i*nhibitors of *<u>a</u>*minoacyl-t*<u>R</u>*NA synthetases), constitute a family of selective inhibitors of distinct types of aaRSs; (ii) the aaRSs encoded within tiarin BGCs represent antibiotic-resistant variants of the target proteins, and, thus, they confer self-immunity to the producer; (iii) the amino acid residue preceding the conserved glutamine in the precursor peptide determines the specificity of the antibiotic to a particular aaRS type. Proving these hypotheses correct would provide a straightforward sequence-based rule for predicting the target specificity of individual tiarins (**Fig. 2a**, column “tiarin specif.”).

### TFX targets arginyl-tRNA synthetase in Rhizobium leguminosarum

In the trifolitoxin precursor, TfxA, an Arg residue precedes the conserved Gln (**Fig. 1e**). According to our hypothesis, TFX should therefore target arginyl-tRNA synthetase (ArgRS). To test this experimentally, we selected the TFX-susceptible *R. leguminosarum* bv. *phaseoli* 4292 (hereafter Rlp_4292) as an indicator strain (**Fig. 1a**). Plasmid-borne copies of *Rlp*_4292 arginyl- or isoleucyl-tRNA synthetase genes (*Rlp-argRS*^HSK^ and *Rlp-ileRS*^HSK^, respectively) were overexpressed in their native host, and the resulting strains were compared for TFX susceptibility using an agar diffusion assay with serial dilutions of the antibiotic. Likely due to antibiotic titration by the overexpressed target enzyme, cells expressing *Rlp-argRS^HSK^* tolerated approximately fourfold higher concentration of TFX than those harboring *Rlp-ileRS^HSK^* or empty-vector control (**Extended Data Fig. 1**). This result is consistent with the prediction that ArgRS may be the cellular target of the antibiotic.

To determine whether TFX selectively inhibits *Rlp*-ArgRS^HSK^, we analyzed changes in tRNA aminoacylation in *Rlp_*4292 upon antibiotic treatment. Total tRNA was extracted from the TFX-treated and untreated cells to estimate levels of aminoacylated tRNAs. An LC-MS analysis revealed an approximately 50-fold decrease in arginyl-tRNA^Arg^ abundance in the TFX-treated cells, whereas the abundance of all other aminoacylated tRNA species remained largely unchanged between the TFX-treated and control cells (**Fig. 3a, Supplementary Fig. 4, Supplementary Data 3**). Together, these results indicate that TFX induces growth arrest by selectively inhibiting tRNA^Arg^ aminoacylation.

**Figure 3.**
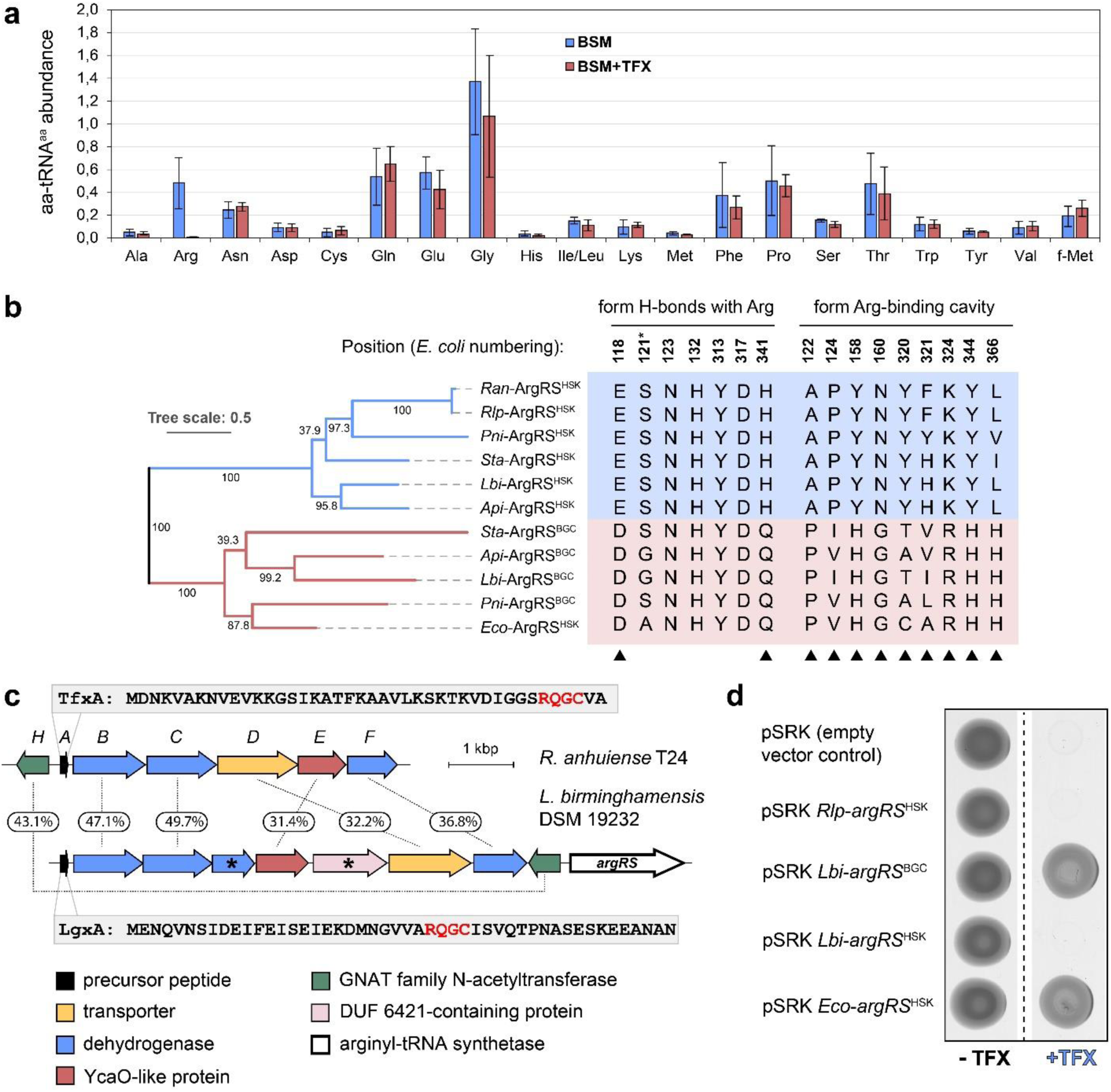
Arginyl-tRNA synthetase is the molecular target of TFX. **a.** Comparison of individual aminoacylated tRNA abundance in *Rlp_*4292 cells treated and untreated with TFX. Mean values from three independent biological replicates were compared using a series of t-tests with Benjamini-Hochberg correction for multiple comparisons^44^. A significant decrease was observed only for only for Arg-tRNA^Arg^ (BH-adjusted p-value = 0.022). **b.** Maximum likelihood phylogenetic tree of ArgRS homologs. Numbers at branching points indicate branch support values from 1000 replicates (SH-like aLRT). Selected columns from the alignment of ArgRS sequences from the genomes of the listed strains are shown on the right. Amino acids forming hydrogen bonds (H-bonds) with arginine or involved in the ligand-binding cavity formation (based on the structure of *Eco-*ArgRS in complex with Arg, PDB ID: 4OBY^18^) are shown. Black arrowheads mark positions where housekeeping and BGC-encoded ArgRSs differ in amino acid identity. Ala121 (marked with an asterisk) forms an H-bond with arginine through its carbonyl; the remaining six residues coordinate the ligand through their side chains. *Ran* – *Rhizobium anhuiense* T24, *Rlp* – *Rhizobium leguminosarum* 4292, *Pni* – *Paenarthrobacter nitroguajacolicus* xwA3 200, *Sta* – *Staphylococcus* sp. MI 10-1553, *Lbi* – *Legionella birminghamensis* DSM 19232, *Api* – *Acinetobacter pittii* DSM 25618, *Eco* – *Escherichia coli* BW25113 **с.** Comparison of the *tfx* and *tfx*-like BGCs from *Legionella birminghamensis* DSM 19232. Predicted functions of the encoded proteins are listed below. Percent sequence identities between homologs are indicated. Genes marked with a black asterisk lack a corresponding homolog in the *tfx* BGC. Amino acid sequences of the precursors are shown for both BGCs, with RQGC motifs in the core peptides highlighted in red. **d.** Growth of the *R. leguminosarum* 4292 control strain and derivatives expressing the indicated aaRSs on media supplemented with 2 µM TFX and on antibiotic-free control plates.

Given that direct interaction between TFX and ArgRS is likely responsible for the observed phenotype, we examined features of the enzyme that could confer resistance to TFX. To this end, we compared sequences of ArgRSs from tiarin BGCs responsible for the biosynthesis of the close structural homologs of TFX (*i.e*., those containing the *ycaO* gene, see **Fig. 1e, Supplementary Data 2**) with the ArgRS^HSK^ copies encoded in the same genomes. ArgRS sequences from *Escherichia coli* BW25113 (*Eco-*ArgRS^HSK^), the TFX-producing strain *R. anhuiense* T24 (*Ran*-ArgRS^HSK^), and the TFX-susceptible *Rlp_*4292 (*Rlp*-ArgRS^HSK^) were also included in the analysis. Close inspection of the ArgRSs sequence alignment **(Supplementary Fig. 5)** revealed that the BGC-associated and housekeeping enzymes differ substantially in conserved amino acids forming the arginine binding pocket^18^ (**Fig. 3b, Extended Data Fig. 2**). Notably, *Eco*-ArgRS^HSK^ clustered together with the BGC-associated ArgRSs, suggesting that these enzymes retain catalytic functionality despite a markedly altered ligand-binding site.

Since the *tfx* BGC from *R. anhuiense* T24 lacks a gene for ArgRS, we chose the tiarin BGC from *Legionella birminghamensis* DSM 19232 (*Lbi*) for experimental validation. This cluster encodes a peptide with an RQGC motif in the core, identical to the tetrapeptide post-translationally modified in TFX, and a set of *tfx*-like biosynthetic genes extended by an additional dehydrogenase (**Fig. 3c**). The BGC also includes a gene of arginyl-tRNA synthetase (*Lbi-argRS*^BGC^), while a second *argRS* copy (*Lbi-argRS*^HSK^) is located in a distant genomic locus. We expected that the product of the *L. birminghamensis* tiarin BGC would be structurally similar to TFX and that *Lbi-*ArgRS^BGC^ would confer protection to *Rlp*_4292 against the antibiotic. We first confirmed that both *Lbi-*ArgRS^BGC^ and *Lbi-*ArgRS^HSK^ can aminoacylate rhizobial tRNA in an *in vitro* assay (**Supplementary Fig. 6**). Next, we tested the ability of *Rlp_*4292 expressing housekeeping- or BGC-associated ArgRSs to grow in the presence of 2 µM TFX (∼100X minimal inhibitory concentration for wild-type *Rlp*_4292). As expected, a high concentration of the antibiotic inhibited the growth of cells overexpressing *Rlp*-ArgRS^HSK^ or *Lbi-*ArgRS^HSK^. In contrast, overexpression of *Lbi-*ArgRS^BGC^ rendered the cells resistant to TFX (**Fig. 3d**). This high level of resistance suggests that TFX cannot bind to *Lbi-*ArgRS^BGC^ and indicates that tiarins from *R. anhuiense* and *L. birminghamensis* share the same molecular target. Furthermore, consistent with our bioinformatic prediction (**Fig. 3b**), overexpression of *Eco*-ArgRS^HSK^ also conferred TFX resistance to *Rlp_*4292 (**Fig. 3d**).

Taken together, our data demonstrate that TFX selectively inhibits the aminoacylation of tRNA^Arg^ and that BGC-encoded variants of ArgRS confer immunity to the antibiotic.

### Rlp-ArgRS binds a proteolytic fragment of TFX

To elucidate the mechanism of ArgRS inhibition by TFX, we first tested the antibiotic effect in an *in vitro* aminoacylation assay using recombinant *Rlp-*ArgRS^HSK^. Strangely, no effect of the antibiotic was detected under these conditions (**Fig. 4a)**. We then asked whether *Rlp-*ArgRS^HSK^ could be isolated from the TFX-exposed cells in complex with the antibiotic. To this end, we overexpressed *Rlp-*ArgRS^HSK^ in the *Rlp_*4292 host and purified it from cells treated with two different concentrations of TFX, along with an untreated control (**Supplementary Fig. 7**). The purified protein samples were then analyzed by mass-spectrometry under native conditions. Strikingly, in addition to the mass-ion corresponding to the ionized *Rlp-*ArgRS^HSK^ (average MW = 67,169.14 Da), a dose-dependent appearance of the mass-ion with a +609 Da mass shift was detected in the samples isolated from the TFX-treated cells (**Fig. 4b**). Moreover, in the low *m/z* range of the same spectra, we observed the [M+H]^+^ ion at *m/z* = 609.256. The observed mass shift is lower than expected for intact TFX but matches the theoretical value for the C-terminal six residues of the dehydrated form of TFX (expected *m/z* = 609.2562 [M+H]^+^) with an accuracy below 2 ppm (**Fig. 4c**). LC-MS(/MS) analysis of the same sample showed that the association was noncovalent, and tandem MS analysis further confirmed that this compound (which we further refer to as TFX*^6C^) originated from TFX (**Extended Data Fig. 3**). These findings indicate that ArgRS interacts not with the intact TFX but with a C-terminal fragment of the antibiotic, suggesting that proteolytic cleavage of the Ser5-Arg6 peptide bond is required for target recognition.

**Figure 4.**
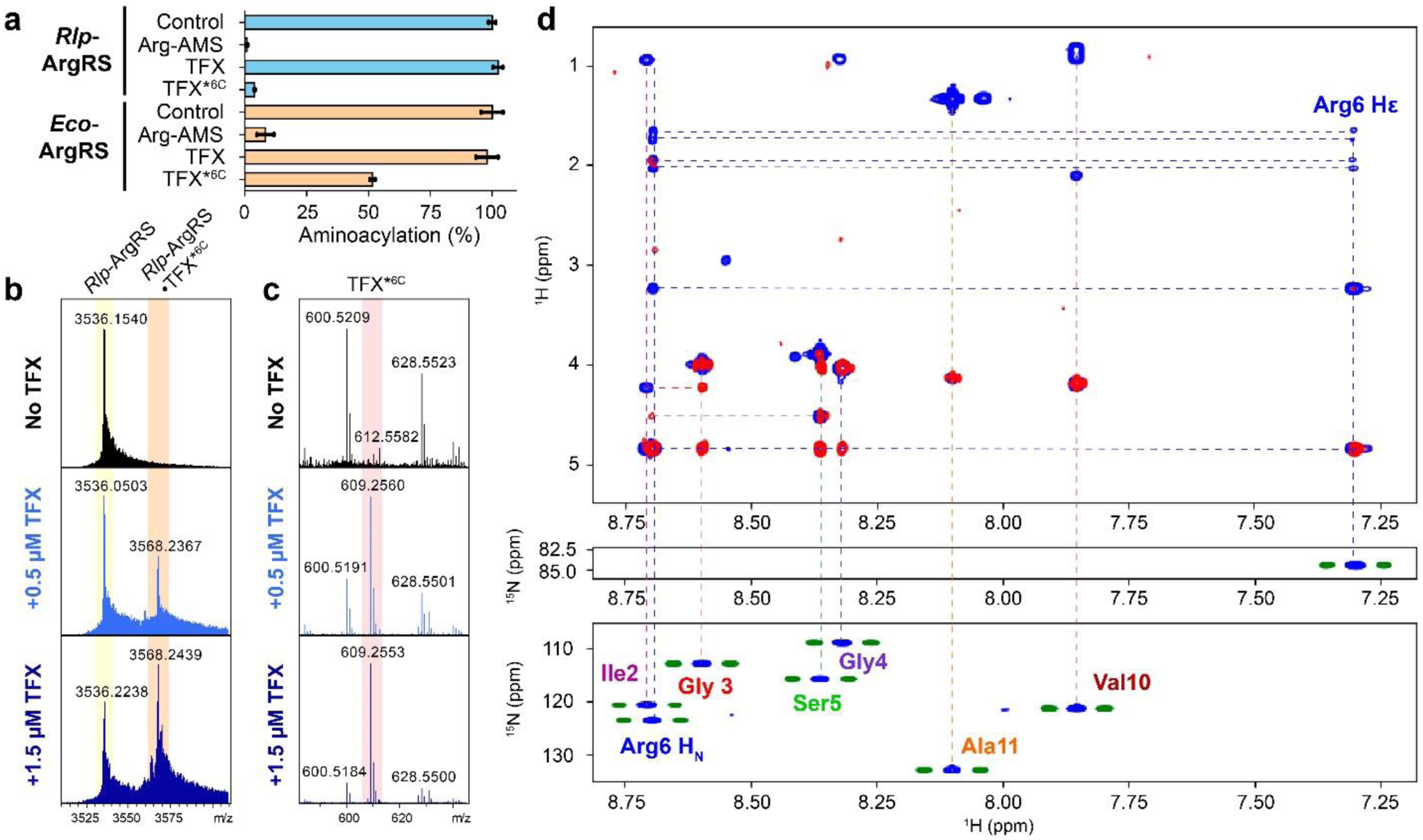
*Rlp*-ArgRS binds a proteolytic fragment of TFX. **a.** Inhibition of *Rlp*- or *Eco*-ArgRS-catalyzed tRNA^Arg^ aminoacylation by TFX or its C-terminal warhead (TFX*^6C^) *in vitro*. 5’-O-(argynilsulfamoyl)adenosine (Arg-AMS) serves as a positive control. **b.** ESI-MS spectra of *Rlp*-ArgRS purified from TFX-treated and control Rlp_4292 cells recorded under native conditions. Note the appearance of additional peaks corresponding to *Rlp*-ArgRS bound to processed TFX (TFX*^6C^) in the samples from TFX-treated cells. **c.** The peaks of free TFX*^6C^ appeared in the low *m/z* region of the spectra of ArgRS samples from TFX-treated cells. **d.** ^1^J ^1^H-^15^N coupling constants for the peptidic part of TFX. (bottom) ^1^H, ^15^N HSQC spectrum of the backbone region of TFX with (blue) or without (green) ^15^N decoupling during the acquisition time. All nitrogens show the characteristic ^1^J value of 90Hz as expected for a directly protonated amide nitrogen. (middle) Cross peak of the Arg6 side chain HN group with (blue) or without (green) ^15^N decoupling during the acquisition time. (top) Planes of the (blue) TOCSY- and (red) NOESY-HSQC spectra with the corresponding assignments.

However, proteolytic processing of TFX conflicts with the TFX structure proposed by Lethbridge *et al*.^8^, in which Arg6 is dehydrated or bears a hydroxyl group attached to its Nα atom (**Fig. 1d**). Either of these modifications would hinder, if not completely prevent, proteolysis between Ser5 and Arg6^19^.

To resolve this discrepancy, we therefore prepared a ^15^N-labeled sample and revisited the structure of TFX by NMR. ^1^H and ^13^C NMR confirmed experimental data of Lethbridge *et al*.^8^ (**Extended Data Table 1**), but did question a modification of Nα or Cα atoms in Arg6. The ^1^H, ^15^N HSQC spectrum showed seven cross peaks with ^15^N chemical shifts in the 110-130ppm range, as expected for regular amide moieties (**Fig. 4d**). No signal corresponding to the NH_2_ group of the modified glutamine was observed. However, a cross peak at 83.6ppm consistent with the ^15^N resonance of an arginine side chain was detected under the same conditions. We assigned all amide peaks, including that of Arg6 (see **Supplementary Discussion** and **Supplementary Fig. 8-11** for detailed description) and recorded the ^1^H, ^15^N HSQC spectrum without ^15^N decoupling during acquisition; the J coupling constant of 90Hz found for seven residues (Arg6 included) confirms the absence of modification of these amide moieties (**Fig. 4d**). Moreover, the Hα proton of Arg6 was visible at resonance at 4.8ppm when spectrum was recorded at 280K in D_2_O (**Extended Data Fig. 4**). In conclusion, whereas our assignments are in excellent agreement with those of Lethbridge *et al.*, we find no evidence for the proposed modification of Arg6. Together, NMR data and the high-resolution MS-derived chemical formula of TFX (**Fig. 1c**) suggest that its fluorophore structure differs from that proposed by Lethbridge *et al*.^8^. While alternative structures compatible with both MS and NMR data can be envisaged (**Supplementary Fig. 12**), resolving this discrepancy will require further investigation. Nonetheless, our results indicate that Ser5 and Arg6 are connected by a conventional peptide bond that can be cleaved by common cellular peptidases.

To confirm that the peptide bond between Ser5 and Arg6 can indeed undergo proteolytic cleavage, we performed *in vitro* processing of TFX using *E. coli* aminopeptidases PepN and PepB. Consistent with our predictions, the resulting proteolytic fragment [M+H]^+^ at *m/z* 609.2532 (**Extended Data Fig. 5**) matched the compound isolated from cells (**Fig. 4b,c**), thus corroborating the NMR data.

Finally, to test whether processed TFX can inactivate *Rlp-*ArgRS^HSK^ *in vitro*, we repeated the *in vitro* aminoacylation assay this time using TFX*^6C^. Unlike TFX, which showed no inhibitory activity, TFX*^6C^ strongly inhibited the aminoacylation of tRNA^Arg^ by *Rlp*-ArgRS^HSK^ **(Fig. 4a)**. Consistent with our *in vivo* data (**Fig. 3d**), inhibition of *Eco*-ArgRS-catalyzed aminoacylation by TFX*^6C^ under the same conditions was substantially weaker (**Fig. 4a)**. Together, our data provide evidence that only the proteolytically processed form of TFX inhibits arginyl-tRNA synthetase.

### TFX acts as a Trojan-horse antibiotic

To further understand the mechanism of TFX action, we performed a genome-wide transposon sequencing (Tn-seq) screen^20^ to identify genes that affect rhizobia survival in the presence of the antibiotic. A library of *Rlp_*4292 mutants containing random single transposon insertions was grown in media supplemented with varying subinhibitory concentrations of TFX. The library strains grown without TFX addition served as the control. The screen revealed dose-dependent changes in Tn insertion frequencies in multiple genes upon TFX treatment (**Supplementary Data 4**). The most prominent enrichment in Tn insertions was observed for the *pepN* gene coding for a Zn^2+^-dependent aminopeptidase and in the genes of the *yejABEF* operon encoding an inner membrane ATP-binding cassette (ABC) oligopeptide transporter, suggesting that inactivation of these genes confers resistance to TFX (**Fig. 5a,b,d, Extended Data Fig. 5**).

**Figure 5.**
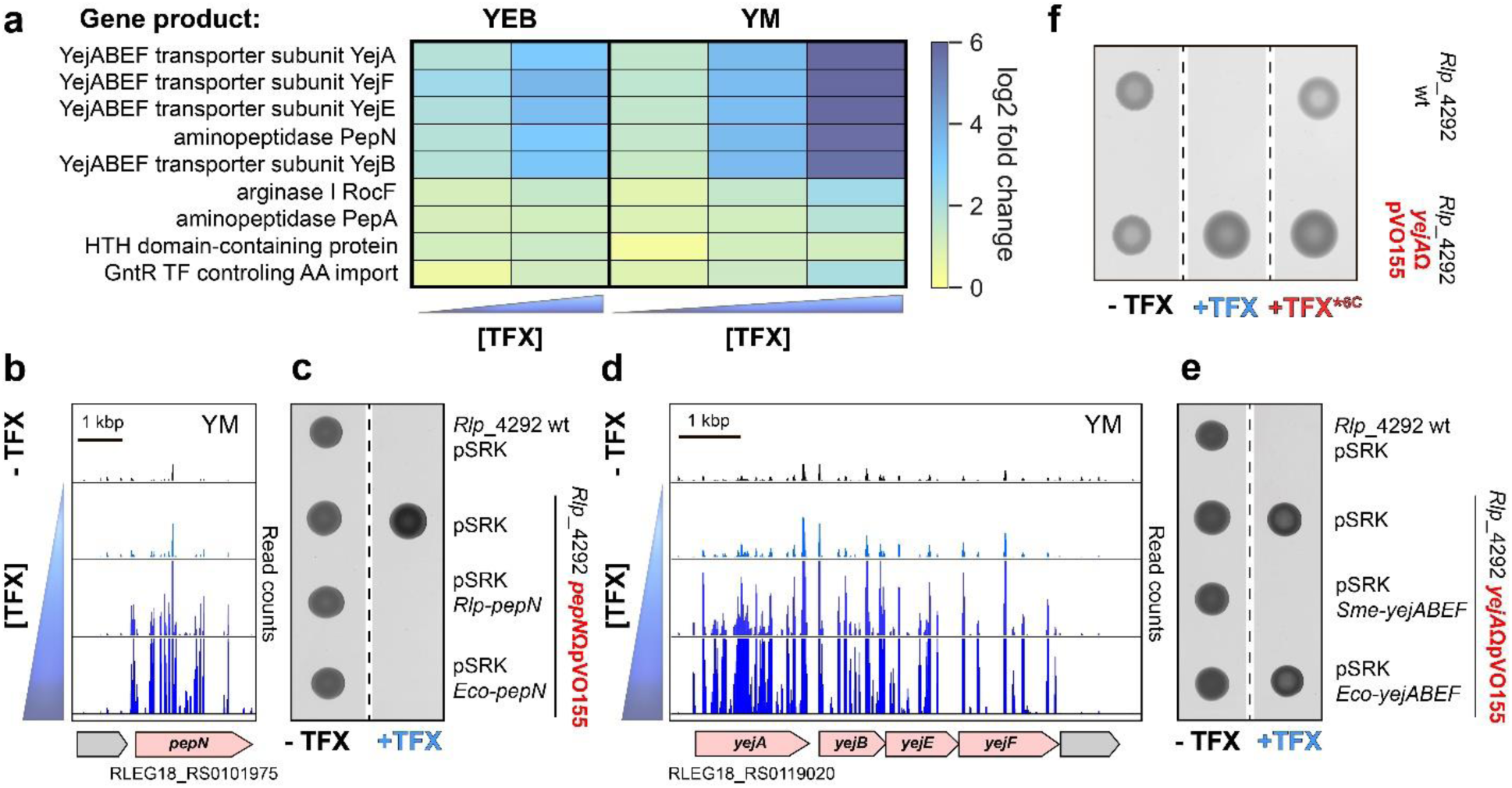
TFX is a Trojan horse antibiotic. **a.** Heat map showing increases in transposon insertion counts for selected genes of *Rlp*_4292 upon TFX treatment in two cultivation media (YEB and YM). **b,d.** Tn-seq results for selected genomic regions of *Rlp*_4292. Histograms indicate the abundance of mutants in the Tn-seq population under the indicated conditions. An increased frequency of insertions in the presence of TFX reflects improved survival of the mutant and therefore the requirement of the corresponding gene product for TFX activity. **c.** Growth of *Rlp*_4292 and derivatives carrying a plasmid insertion–inactivated *pepN* on media with and without TFX. **d.** Growth of *Rlp*_4292 and derivatives carrying a plasmid insertion–inactivated *yejA* on media with and without TFX. **f.** Growth of wild-type *Rlp*_4292 and its derivative with impaired TFX import on media without TFX, with 0.5 µM TFX, and with 0.5 µM *in vitro* processed TFX (TFX*^6C^).

Increased TFX resistance of *Rlp*_4292 as a result of inactivation of a broad-specificity aminopeptidase PepN supports our finding that proteolytic cleavage is required for TFX activation (**Fig. 4a**). In addition to *pepN*, Tn insertions in a gene of another aminopeptidase, PepA, were also enriched (**Fig. 5a**), although to a much lesser extent than for PepN. To validate the primary role of PepN in TFX processing, we generated the *Rlp*_4292 *pepN*ΩpVO155 mutant and tested its susceptibility to TFX. Consistent with the proposed role of PepN in TFX processing, inactivation of its gene rendered *Rlp*_4292 resistant to the antibiotic, whereas the complementation of *Rlp*_4292 *pepN*ΩpVO155 with a plasmid-borne copy of *Rlp*-*pepN* or its *E. coli* homolog (*Eco*-*pepN*) restored susceptibility (**Fig. 5c**).

The abundant transposon insertions in the *yejABEF* operon (**Fig. 5d**) suggest that, like some other peptide antibiotics^21,22^, TFX exploits this oligopeptide transporter to reach its intracellular target. Consistent with this conjecture, the *Rlp_*4292 strain with *the yejA* gene inactivated by a plasmid insertion (*Rlp_*4292 *yejA*ΩpVO155) was resistant to high concentrations of TFX (**Fig. 5e**), indicating the primary role of the YejABEF transporter in the TFX uptake. Overexpression of the *yejABEF* operon of a phylogenetically close *S. meliloti* compensated for the loss of Yej transporter function in *Rlp_*4292 *yejA*ΩpVO155 (**Fig. 5e)**. In contrast, overexpression of *E. coli yejABEF* did not restore *Rlp*_4292 susceptibility to trifolitoxin (**Fig. 5e**). Thus, along with the *Eco-*ArgRS^HSK^-mediated antibiotic resistance (**Fig. 3d**), impaired antibiotic uptake can contribute to the insensitivity of *E. coli* to TFX^23^.

To examine the role of the TFX N-terminus in its biological activity, we tested the susceptibility of *Rlp_*4292 to the *in vitro*-prepared TFX warhead (TFX*^6C^) (**Extended Data Fig. 6**). While the growth of *Rlp_*4292 in the presence of TFX was inhibited, it grew normally in the presence of the TFX*^6C^ (**Fig. 5f**). These data show that while the unmodified N-terminal part of TFX directly interferes with ArgRS inhibition, it is essential for efficient TFX uptake by susceptible rhizobia.

Altogether, our data show that TFX acts as a Trojan horse antibiotic: (1) its N-terminal pentapeptide serves as a bait and mediates uptake of the antibiotic *via* the YejABEF oligopeptide transporter; (2) once TFX is in the cytoplasm, subsequent intracellular proteolysis by non-specific aminopeptidases, primarily PepN, leads to the release of the post-translationally modified TFX*^6C^ warhead that targets ArgRS.

### TfxH confers immunity to the producer by acetylating processed TFX

Unlike many tiarin BGCs, the *tfx* BGC does not encode a resistant variant of aaRS, which could protect the host strain from self-intoxication. Earlier studies assigned an immunity role to *tfxG*, encoding a putative phosphotransferase, or to *tfxE*^10^, encoding a YcaO-like enzyme^24^ whose biochemical function was unknown at the time. Bioinformatics analysis identified an additional gene, *tfxH,* encoding a GCN5-related N-acetyltransferase, which is located adjacent to the *tfx* BGC (**Fig. 2b, Supplementary Data 2**). We speculated that tiarin BGC-associated GNATs could serve as self-immunity determinants. To identify which gene(s) of the *tfx* BGC could confer protection against TFX, we tested whether expression of *tfxE, tfxG,* and *tfxH* from a plasmid would render TFX-susceptible *Rlp_4292* resistant to the antibiotic. Since active export of a toxic compound from cells is a common mechanism of resistance, a plasmid expressing *tfxD*, a <u>m</u>ultidrug <u>a</u>nd toxic compound <u>e</u>xtrusion (MATE) family transporter, was used as a positive control. Neither *tfxE* nor *tfxG* conferred noticeable protection against TFX (**Fig. 6a**). In contrast, expression of *tfxH,* as well as *tfxD*, enabled the growth of *Rlp_*4292 on medium containing 0.2 µM TFX (**Fig. 6a**). To elucidate the biochemical basis of resistance to tiarins provided by acetyltransferases, we combined the purified TfxH and acetyl coenzyme A (AcCoA) in an *in vitro* reaction with TFX. However, MALDI-TOF MS analysis of the reaction products revealed no product with the mass expected for the acetylated TFX (**Fig. 6b**), suggesting that TfxH cannot acetylate mature TFX under the tested conditions. We then hypothesized that TfxH might instead recognize the proteolytically processed TFX, which can occasionally form in the producer due to nonspecific degradation by cellular peptidases. We therefore repeated the experiment, this time including the *E. coli* aminopeptidases PepB and PepN in the reaction mixtures. As expected, aminopeptidase treatment removed five N-terminal residues, resulting in the appearance of an [M+H]^+^ ion at *m/z* = 609.3, which corresponds to TFX*^6C^ (**Fig. 6c**) and when TfxH and AcCoA were present in the reaction, a new [M+H]^+^ mass ion at *m/z* = 651.3 appeared, consistent with acetylated TFX*^6C^ (**Fig. 6c**). The acetyl moiety is most likely attached to the α-amino group of the Arg residue, which explains the previously observed lack of modification of the unprocessed TFX.

**Figure 6.**
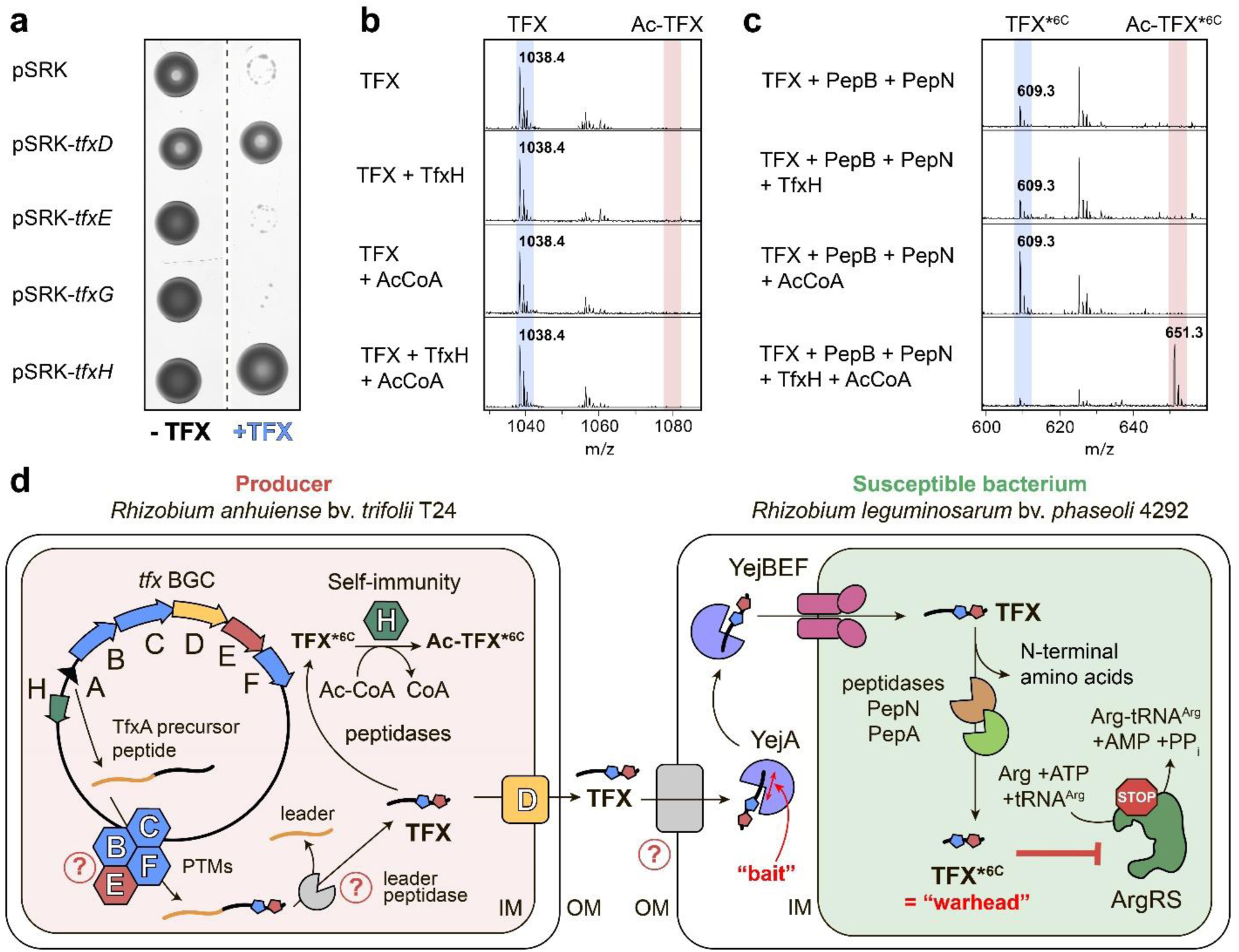
TFX self-resistance in the producer and an overview of the TFX mode of action. **a.** Growth of wild-type *R. leguminosarum* 4292 and its derivatives expressing individual *tfx* BGC genes on media with 0.2 µM TFX and on TFX-free control media. **b.** MALDI-TOF MS analysis of *in vitro* reactions of TFX (*m/z* = 1038.4 [M+H]^+^) modification by TfxH. AcCoA – acetyl coenzyme A. **c.** MALDI-TOF MS analysis of TfxH-mediated modification of TFX pretreated with PepB and PepN. Mass ion at *m/z* = 609.3 [M+H]^+^ corresponds to the N-terminally processed TFX (TFX*^6C^), [M+H]^+^ at *m/z* = 651.3 [M+H]^+^ – to acetylated TFX*^6C^ (Ac-TFX*^6C^). **d.** Proposed scheme outlining TFX biosynthesis and its mode of action. The blue fluorescent chromophore and the thiazoline in TFX are shown as blue and red pentagons, respectively. The processes, for which the proteins involved or the detailed mechanisms remain unknown, are denoted with red question marks. ArgRS – arginyl-tRNA synthetase, TFX*^6C^ – N-terminally processed TFX, which can be modified by TfxH GNAT and binds to *Rlp*-ArgRS^HSK^, IM – inner membrane, OM – outer membrane, PTMs – posttranslational modifications.

Together, these results indicate that the TfxH N-acetyltransferase functions as a second line of defense, complementing the TFX transporter TfxD. TfxH detoxifies the TFX warhead, if it is generated in the producer by endogenous peptidases, thereby protecting the housekeeping arginyl-tRNA synthetase and helping the host avoid suicide.

## DISCUSSION

Trifolitoxin (TFX) inhibits the growth of susceptible rhizobia at nanomolar concentrations (**Extended Data Fig. 1**), which places it among the most potent natural antibiotics. Here we show that TFX functions as a Trojan-horse inhibitor: it hijacks the oligopeptide importer YejABEF, which apparently recognizes the unmodified N-terminal portion of TFX as a "bait" and internalizes the molecule. Once inside the cell, broad-specificity aminopeptidases, primarily PepN, remove the N-terminal residues of TFX to release the C-terminal "warhead," a six-residue fragment bearing post-translational modifications. This "warhead" then binds its intracellular target, ArgRS, thereby arresting translation (**Fig. 6d)**.

Our work reveals multiple strategies by which bacteria gain resistance to TFX. We identify N-acetyltransferases as a prevalent mechanism of immunity to TFX and other tiarins. This mechanism mirrors the protective acetylation of the α-amino group of the aminoacyl moiety in isoasparaginyl-adenylate observed for microcin C, another RiPP that targets aaRSs^25^. In addition, expression of a resistant target protein provides substantial protection. We show that some bacteria, such as *E. coli*, harbor housekeeping ArgRS variants intrinsically resistant to TFX (**Fig. 3d**). However, it is worth noting that this type of resistance does not warrant protection against other tiarins that target distinct aaRSs. Differences in YejABEF transporter substrate selectivity are known to determine the activity of many peptide antibiotics^21,22^. It is tempting to speculate that, as with other antibiotics that use peptide transporters^26,27^, altering the length or sequence of the unmodified N-terminal part may allow rational tuning of tiarin’s antibacterial spectrum.

To our knowledge, TFX is the first naturally produced antibiotic that selectively inhibits arginyl-tRNA synthetase^28^. However, the detailed mechanism of its binding to ArgRS, and even its structure, remains elusive. Our data suggest that the previously proposed architecture of the glutamine-derived fluorophore^8^ requires revision. Such ambiguity is not uncommon among RiPPs, as revealing the structures of heavily modified peptides is notoriously challenging using mass spectrometry and NMR spectroscopy alone^29,30^. A definitive structural assignment requires further research, which would benefit from orthogonal methods such as stepwise biosynthetic pathway reconstitution and crystallographic or cryogenic electron diffraction analysis.

The characteristic fluorophore of TFX is likely a conserved modification across the tiarin family. This shared scaffold is further tailored by secondary modifications and coupled to an upstream amino acid that directs target recognition, thereby ensuring specific interaction with a particular aaRS. The use of a common scaffold for targeting different families of aaRSs makes tiarins unique among natural antibiotics. Our bioinformatic analysis uncovers a broad diversity of tiarin BGCs distributed across bacterial and archaeal genomes, whose predicted products likely target at least seven class I and at least one class II aaRSs (**Extended Data Fig. 7**). As genomic datasets continue to expand, additional tiarins with novel specificities will almost certainly emerge. The essential role of aaRSs and the presence of 20 potential targets in each cell offer new opportunities to expand the spectrum of druggable targets in bacteria and to design next-generation compounds based on this conserved scaffold, possibly even expanding beyond the aaRSs targeted by natural compounds.

## MATERIALS AND METHODS

### Genome mining for tfx-like BGCs across bacterial genomes

Translations of all coding sequences annotated in prokaryotic genomes from the RefSeq database^31^ were downloaded on March 1, 2022, from the NCBI ftp site (https://ftp.ncbi.nlm.nih.gov/). These sequences were used as a database for BLASTP^32^ search with the amino acid sequence of TfxC (NCBI Protein database accession number “WP_183844980.1”) as a query and an E-value cut-off of 10.

Genomic records containing the genes retrieved by the search were downloaded in GenBank file format, and regions including 10 kb upstream and 10 kb downstream from *tfxC* homologs were excised. Domain architectures of all proteins encoded within these regions were annotated using the batch CD-search web tool (https://www.ncbi.nlm.nih.gov/Structure/bwrpsb/bwrpsb.cgi)^15^.

BLAST hits were clustered using MMseqs2^12^ with the 95% identity cutoff, and representative sequences from each cluster were used in further analysis. If a representative sequence appeared to result from the translation of a pseudogene or a *tfxC* homolog at the contig end, it was replaced with another protein from the corresponding cluster when possible or excluded from the analysis. A similarity network of representative sequences was constructed with EFI-EST^33^ with an alignment score cut-off of 32 and visualized using Cytoscape^34^.

TfxB (accession number “WP_183844979.1”) homologs were retrieved in another BLASTP search with the same parameters as in the initial TfxC search. Since TfxB and TfxB are homologous, proteins that were identified in both searches were removed from the network if the E-value obtained in the TfxB search was lower than in the TfxC search, and BLAST alignment was attributed to the same domain according to CD-search annotation. If the genes of TfxC and TfxB homologs were present within the same contig, the distances between them were calculated. Connected components containing TfxC homologs encoded within ±2.7 kb from the gene of TfxB homolog were selected for further analysis.

Domains identified by CD-search that correspond to sequence regions aligned by BLAST were excised together with 10 additional amino acids at each side. Sequences shorter than 100 amino acids were excluded from the dataset.

The resulting 79 sequences were combined with five SagB homologs randomly selected from the stand-alone proteins in the TfxC sequence similarity network (**Supplementary Figure 2**), which served as an outgroup for the phylogenetic tree rooting. The alignment was obtained using MUSCLE^35^ with a gap opening penalty of -10 and subsequently trimmed using ClipKIT^36^ with default parameters. The phylogenetic tree was reconstructed using RaxML^37^ with the LG substitution model and gamma-distributed evolutionary rates. The rapid bootstrap procedure converged after 500 iterations according to the extended majority-rule consensus tree (MRE) criterion. The resulting tree was visualized with iTOL^38^.

Selected biosynthetic gene clusters containing TfxC and TfxB homologs were examined manually for the presence of open reading frames coding for putative precursor peptides and genes potentially involved in RiPP posttranslational modification and export. Motif discovery in precursor peptides was performed using MEME^14^ with one expected occurrence of motif per sequence and a minimal motif length of four amino acid residues. All proteins encoded in the genomes harboring *tfx*-like BGCs were scanned with hidden Markov models of motifs specific to aminoacyl-tRNA synthetases using the tool available at the prokaryotic AARS database website (https://bioinf.bio.uth.gr/aars/)^39^. A protein was identified as an aaRS if it contained at least two motifs. Predicted precursor peptides were aligned using MUSCLE^35^ with default parameters.

### Bacterial strains and growth conditions

Bacterial strains used in the study are listed in **Supplementary Table 2**. *Rhizobium* strains were cultivated in YM medium (10 g L^-1^ mannitol, 0.5 g L K_2_HPO_4_, 0.2 g L^-1^ MgSO_4_, 0.1 g L^-1^ NaCl, 1 g L^-1^ yeast extract, pH 6.8) or modified BSM medium^40^ (10 g L^-1^ mannitol, 1.21 g L^-1^ glutamic acid monosodium salt monohydrate, 0.179 g L^-1^ Na_2_HPO_4_, 0.05 g L^-1^ MgSO_4_, 0.04 g L^-1^ CaCl_2_, 0.02 g L^-1^ FeCl_3_, 1 mg L^-1^ thiamine, 150 µg/L biotin, pH 6.8). *E. coli* strains were grown in LB medium (10 g L^-1^ tryptone, 5 g L^-1^ yeast extract, 5 g L^-1^ NaCl, pH 7.0) or 2xYT medium (16 g L^-1^ tryptone, 10 g L^-1^ yeast extract, 5 g L^-1^ NaCl, pH 7.0). Rhizobia and *E. coli* were cultivated at 28 °C and 37 °C, respectively. Where appropriate, antibiotics were used at the following concentrations: ampicillin (Ap), 100 µg·mL^-1^; kanamycin (Km), 100 µg·mL^-1^; rifampicin (Rf) 40 µg·mL^-1^; chloramphenicol (Cm), 34 µg·mL^-1^; gentamycin (Gm), 50 µg·mL^-1^.

For rhizobial growth inhibition assays, 5 µL of the indicated overnight cultures were spotted onto BSM agar plates supplemented with the specified concentrations of TFX. Plates were incubated for 48 h before imaging.

For agar diffusion assays, soft BSM agar was inoculated with the indicated rhizobial strain to prepare a uniform lawn. Then, 3-µL drops of 2-fold serial dilutions of TFX were spotted onto the agar surface. Plates were incubated for 48 h to allow lawn formation and growth inhibition zones development.

### Molecular cloning

Vectors and oligonucleotide primers used in this study are listed in **Supplementary Tables 2 and 3**, respectively. The genes *tfxH*, *tfxD*, *tfxE*, and *tfxG* were PCR-amplified using *R. anhuiense* T24 genomic DNA as the template, and *Rlpl-pepN*, *Rlp-argRS*^HSK^, and *Rlp-ileRS*^HSK^ were amplified from *R. leguminosarum* bv. *phaseoli* 4292 genomic DNA. Lbi-argRS^BGC^ and Lbi-argRS^HSK^, based on the *Legionella birminghamensis* DSM 19232 genome, were obtained as synthetic DNA fragments (Twist Bioscience) (**Supplementary Table 4**). *Eco-pepN* and *Eco-argRS*^HSK^ were amplified using *E. coli* BW25113 genomic DNA.

Gene fragments were assembled into the multiple cloning sites of the corresponding vectors listed in **Supplementary Table 2** using a standard restriction–ligation protocol. The pSRK-T5-lac vector was constructed by inserting a PCR-amplified fragment of the pCA24N vector^41^ into the pSRK backbone. Sequences encoding Strep and 6×His tags were added to *Rlp-argRS*^HSK^ by PCR before cloning into the pSRK-T5-lac vector.

### Protein expression and purification

An overnight culture of *E. coli* BL21(DE3) cells transformed with the corresponding vectors (**Supplementary Table 2**) was diluted 100-fold into 2×YT medium and grown at 37 °C until the OD₆₀₀ reached 0.6. Protein expression was induced with 0.2 mM IPTG, and cells were incubated at 25 °C for 16 h. Cells were harvested by centrifugation, resuspended in Lysis Buffer (20 mM Tris-HCl pH 7.5, 150 mM NaCl, 5 mM imidazole, 2 mM β-mercaptoethanol, 1 mM PMSF), lysed by sonication, and clarified by centrifugation at 16,000 × g for 20 min at 4 °C. The cleared lysate was applied to a 1-mL HiTrap TALON column (GE Healthcare Bio-Sciences) pre-equilibrated with Lysis Buffer. The column was washed with 25 bed volumes of Wash Buffer (20 mM Tris-HCl pH 7.5, 150 mM NaCl, 5 mM imidazole, 2 mM β-mercaptoethanol), and bound protein was eluted with 5 mL of Elution Buffer (20 mM Tris-HCl pH 7.5, 150 mM NaCl, 300 mM imidazole). Collected fractions were buffer-exchanged into Storage Buffer (20 mM Tris-HCl pH 7.5, 200 mM NaCl, 1 mM EDTA, 10% v/v glycerol) using 10-kDa cutoff dialysis tubing. Proteins were stored at –80 °C until further use.

For purification of *Rlp*-ArgRS in complex with processed TFX, *Rlp*_4292 cells were transformed with the pSRK-pT5-lac_*Rlp-*ArgRS^HSK^ expression vector. One liter of BSM medium supplemented with 1% (w/v) yeast extract was inoculated with 20 mL of *Rlp*_4292/pT5-lac-*Rlp-*ArgRS^HSK^ starter culture. Protein expression was induced with 1 mM IPTG when the culture reached an OD_600_ of 0.8. Following induction, cells were incubated at 20 °C for 16 h, supplemented with 2 µM TFX, and further incubated at 20 °C for 6 h. Cells were harvested and resuspended in the TSBB buffer (20 mM Tris-HCl pH 8.0, 300 mM NaCl, 2 mM β-mercaptoethanol) supplemented with 1 mg·mL⁻¹ lysozyme and incubated on ice for 30 min. Triton X-100 was then added to a final concentration of 1% before sonication. The clarified lysate was applied to a 1 mL Strep-Tactin superflow column (IBA Lifesciences). After washing with the TSBB buffer supplemented with 1 mM MgSO_4_, the protein was eluted with the same buffer additionally supplemented with 2.5 mM desthiobiotin, then buffer exchanged to After buffer exchange to 10 mM MS grade ammonium acetate pH 7.0 by ultrafiltration at 30 kDa cut-off on Amicon 0.5 mL cartridges and subjected to MS analysis. Protein purity was assessed by SDS–PAGE.

### Purification of TFX and TFX*^6C^

An overnight culture (40 mL) of *R. anhuiense* T24 was inoculated into 2 L of BSM medium. For NMR analysis, *R. anhuiense* T24 was grown in BSM supplemented with [^15^N]-glutamate. Isotopically labeled glutamate was synthesized from [^15^N]-ammonium chloride and α-ketoglutarate as described in ref.^42^. Cultures were grown with shaking at 28 °C for ∼30 h. Cells were pelleted (14,000 x *g*, 30 min, RT), and the supernatant was collected, filtered through a 0.22-µm nitrocellulose filter, and applied to a 10 g HF C18 Bond Elut cartridge (Agilent). The cartridge was washed sequentially with 150 mL of 5 mM K-phosphate buffer pH 5.8, and 100 mL of 5% acetonitrile in the same buffer. Bound peptides were eluted with 50 mL of 15% acetonitrile in 5 mM K-phosphate buffer pH 5.8, vacuum-dried, and resuspended in 1 mL of deionized water. A 100-µL aliquot of this sample was subjected to reverse-phase HPLC on an Agilent ZORBAX Eclipse XDB-C18 column (4.6 x 250 mm, 5 µm particle size). The column was pre-equilibrated with 5 mM K-phosphate buffer pH 5.8. Peptides were separated using a 5–17% linear gradient of acetonitrile at a flow rate of 1 mL min^-1^ (**Supplementary Fig. 13a**). Peaks were monitored at 302 nm, the characteristic absorbance of the blue-fluorescent TFX chromophore^8^. Collected fractions were vacuum-dried, redissolved in deionized water, and subjected to a second round of purification on the same column pre-equilibrated with 20 mM triethylammonium acetate (TEAA) pH 7.0 (**Supplementary Fig. 13b**). Purified TFX was eluted using a 7–17% linear gradient of acetonitrile at 1 mL min^-1^, with detection at 302 nm. Fractions were vacuum-dried, resuspended in deionized water, and analyzed by MALDI-TOF MS and rhizobial growth-inhibition assays as described above. Purified TFX was stored at –20 °C.

Purified TFX (300 µM) was incubated with 3 µM *E. coli* PepN and 5 µM PepB aminopeptidases in 20 mM Tris-HCl, pH 7.0, containing 20 mM NaCl and 0.2 mM MnCl₂ at 30 °C for 40 min. The reaction mixture was purified by HPLC on the ZORBAX Eclipse XDB-C18 column (4.6 x 250 mm, 5 µm particle size) pre-equilibrated with 0.1 M TEAA pH 6.5. TFX*^6C^ was eluted using a 0–18% linear gradient of acetonitrile, with detection at 302 nm. (**Supplementary Fig. 13c**). Fractions were analyzed by MALDI-TOF MS. Purified TFX*^6C^ was dissolved in deionized water and stored at-20 °C.

### Analysis of rhizobial tRNA aminoacylation abundance in rhizobial cells

Analysis of rhizobial tRNA aminoacylation was performed using three biological replicates. An overnight culture of Rlp_4292 was diluted 10-fold into BSM medium supplemented with 0.2% casamino acids and grown with agitation at 28 °C for 5 h. One half of the culture was then supplemented with 0.02 µM TFX, while the other half served as an untreated control. After 3 h of incubation, cells were harvested by centrifugation at 13,000 x *g* for 10 min at 4 °C. tRNA extraction and acetylation were performed as described previosly^43^.

For aminoacylation efficiency measurement, 20 µg of total rhizobial tRNA fraction was deacylated in 10 mM Tris HCl buffer, pH 9.0, at 30 °C for 30 min, combined with 0.5 µM arginyl-tRNA synthetases and 10 µM arginine in 60 mM HEPES buffer pH 7.6, containing 30 mM KCl, 10 mM MgCl₂, 2.5 mM ATP. Reactions were carried out at 37 °C for 10 min. Products of the *in vitro* aminoacylation reaction were acetylated as described in ref ^43^.

Prior to LC-MS analysis, a total of 10 µg of acetylated aminoacyl-tRNA was digested with 100 U of RNase I (Ambion) and 1,000 U of RNase T1 (Thermo) in 30 mM ammonium acetate pH 7.0 at 37 °C for 30 min.

### Mass-spectrometry

MALDI-TOF spectra were recorded using the UltrafleXtreme II MALDI-TOF-TOF (Bruker), equipped with a neodymium laser (355 nm). Samples were analyzed in the matrix solution containing 40 mg mL^-1^ 2,5-dihydroxybenzoic acid (Sigma) and 0.5% trifluoroacetic acid (TFA) in 30 % acetonitrile. The mass spectra recorded in reflector mode with the measurement accuracy within 0.1 Da. For the fragmentation spectra LIFT mode was used, the accuracy for the detected product ions within the 1 Da range. FlexAnalysis 3.2 software (Bruker) was used for mass-spectra processing.

The high-resolution spectra were recorded as part of the LC-MS analysis on an Agilent 1200 HPLC equipped with the UV and 6550 iFunnel QTOF LC-MS detectors and Jet Stream Technology ion source (Agilent). The analyzed compounds were loaded on the Poroshell 120 SB-C18 column (2.7 µ, 2.1 ⊆ 100 mm, Agilent) at 40 °C using a linear (0 to 80%) gradient of acetonitrile in 5 mM ammonium acetate buffer (pH 5.2) at a 0.2 mL min^-1^ flow rate. The electrospray source was set to positive ion mode at 4 kV, 290 K. Data were analyzed using MassHunter Qualitative Analysis 10.00 software (Agilent).

LC-MS analysis of acetylated aminoacyl adenosines was carried out using an Agilent 1200 HPLC system equipped with UV detection and a 6550 iFunnel QTOF LC-MS with a Jet Stream Technology ion source. RNase digestion products were separated on an Agilent InfinityLab Poroshell 120 SB-C18 column (2.1 × 100 mm, 2.7 µm particle size) at 40 °C using a 0–80% linear gradient of acetonitrile in 5 mM ammonium acetate buffer pH 5.2 at a flow rate of 0.2 mL min^-1^. The electrospray source was operated in positive-ion mode at 4 kV and 290 K. Data were acquired over an *m/z* range of 200–1100 at 2 spectra/s, and fragmentation spectra were collected in AutoMSMS mode. Data analysis was performed using MassHunter Qualitative Analysis 10.00 software (Agilent) with Quartic/Quintic Savitzky–Golay smoothing (30-point window) and the ChemStation integrator (default settings with advanced baseline correction). The areas of the [M+H]⁺ peaks corresponding to the respective aminoacyl-adenosines were divided by the area of the [M+H]⁺ peak of GMP from the same sample to calculate the normalized aminoacyl-tRNA abundances. Means values from three independent biological replicates were compared using a series of 19 t-tests. To account for multiple comparisons, Benjamini-Hochberg procedure was used^44^.

*Rlp*-ArgRS^HSK^ in complex with TFX*^6C^ was analyzed by flow injection in 10 mM ammonium acetate 10 mM pH 7, on an Ultimate 3000 RSLC (Thermo Scientific) connected to a high-resolution ESI-Q-TOF mass spectrometer (Maxis II ETD, Bruker) in positive ion mode on the range *m/z* 500-4000. The same samples were also analyzed on the same instrument by UHPLC-MS using an Acquity Premier Protein BEH C4 column (Waters, 300 Å, 1.7 µm, 2.1 x 150 mm), at a flow rate of 300 μL/min using a 2 to 80 % gradient of solvent B in 15 min, with solvent A = ultra-pure water/0.1% formic acid and solvent B = HPLC-MS grade acetonitrile/0.08% formic acid. The MS detection was done in positive ion mode in the range *m/z* 250-2500. In addition, data-dependent LC-MS/MS was performed in the range *m/z* 50-1000 by collision-induced dissociation, using an excitation voltage of 40 V.

### In vitro aminoacylation inhibition assay

tRNA^Arg^ aminoacylation reaction mixtures containing 10 µM of deacylated *E. coli* tRNA^Arg^, 7.5 µM [^14^C]-arginine (300 mCi/mmol; American Radiolabeled Chemicals), 30 mM KCl, 60 mM HEPES pH 7.6, 10 mM MgCl₂, 2.5 mM ATP, and 0.5 mg/mL bovine serum albumin were supplemented with 50 µM TFX, 50 µM TFX*^6C^, 50 µM 5’-O-(arginylsulfamoyl)-adenosine (Arg-AMS, MedChemExpress), or no inhibitor. Aminoacylation was initiated by the addition of either *Rlp*-ArgRS or *Eco*-ArgRS to a final concentration of 0.2 µM, followed by incubation for 10 min at 28 °C. A reaction lacking an enzyme was included to control background noise. All reactions were performed in triplicate.

From each reaction, 7.5 µL was spotted onto Whatman 3MM filter discs (Cytiva) prerinsed with 25 µL of 10% trichloroacetic acid (TCA). Discs were immediately transferred into ice-cold 10% TCA, washed three times with 200 mL of cold 10% TCA, rinsed once with 100% acetone, and air-dried for 20 min. Dried discs were placed into scintillation vials containing 5 mL of Ultima Gold liquid scintillation fluid (PerkinElmer), and the radioactivity associated with precipitated tRNA was measured using a Hidex 300 SL automatic liquid scintillation counter (2 min per sample).

### In vitro acetylation of TFX

Reactions were carried out at 28 °C for 40 min in 20 mM Tris-HCl pH 7.0 containing 20 mM NaCl, 0.2 mM MnCl₂, 150 µM Ac-CoA, 1 µM TfxH, and 30 µM TFX. Control reactions lacked TfxH, Ac-CoA, or both. For acetylation of TFX*^6C^, the reaction mixture was additionally supplemented with 3 µM PepB and 5 µM PepN. Reaction products were analyzed by MALDI ToF MS.

### Tn-seq

A *Himar1* transposon mutant library of strain *Rlp_*4292 was generated following a previously described method^18^. Mutants were selected on Modified Arabinose Gluconate medium (MAG: 1.3 g*L^-1^ HEPES, 1.1 g L^-1^ MES, 1.0 g yeast extract, 1.0 g L^-1^ L-arabinose,

1.0 g L^-1^ D-gluconic acid, 0.22 g L^-1^ KH_2_PO_4_, 0.25 g L^-1^ Na_2_SO_4_, 0.32 g L^-1^ NH_4_Cl, 6.7 mg L^-1^ FeCl_3_, 15 mg L^-1^ CaCl_2_x2H_2_O, 180 mg L^-1^ MgSO_4_x7H_2_O, 15 g L^-1^ agar, pH 6.6) supplemented with Rif and Kan, on which the otherwise mucoid strain *Rlp*_4292 forms compact colonies. The library consisted of 15x10^6^ independent mutant clones.

Tn-seq screening of strain *Rlp_*4292 in the presence of TFX was performed in two different media, YM and Yeast Extract Beef broth (YEB: 5 g L^-1^ peptone, 5 g L^-1^ beef extract, 5 g L^-1^ sucrose, 1 g L^-1^ yeast extract, 0.4 g L^-1^ MgSO_4_x7H_2_O, pH 7.5), both allowing growth of strain *Rlp_*4292 to high densities, up to OD_600_=3-4 or 8 generations starting from an OD_600_=0.01, which is needed for efficient Tn-seq screening. TFX Minimal Inhibitory Concentrations (MIC) of strain *Rlp_*4292 were found to be 10.5 nM in YM and 168 nM in YEB. An aliquot of the transposon mutant library was diluted in YM or YEB to an OD_600_=0.01 in 25-mL cultures. TFX was added to final concentrations of ½, ¼, or ⅛ MIC. Cultures, including control cultures without TFX, were grown for 36h until reaching an OD_600_=0.25 to 3 (5 to 8 generations of growth). In order to produce Illumina sequencing libraries of the transposon borders, the genomic DNA was extracted and processed as described previously^20^.

The Tn-seq samples were sequenced using an Illumina NextSeq 2000 instrument. The generated data were demultiplexed, trimmed, and mapped to the reference genome of *Rlp_*4292 (GenBank accession no. NZ_KB905373 (chromosome), NZ_KB905374 (plasmid 1), NZ_KB905375 (plasmid 2), NZ_KB905376 (plasmid 3), and NZ_KB905377 (plasmid 4) as described before^20^. FeatureCounts^45^ was used to evaluate the number of reads by gene.

To identify the genes with enhanced fitness under the TFX selection conditions, the total read count per gene was normalized to both the number of TA dinucleotides within that gene (Himar1 transposons insert exclusively at TA dinucleotides^46^) and the total read count for the corresponding sample. The 20% of genes having the lowest normalized values in the control sample were excluded from further analysis due to insufficient statistical power. Differences in the log₂-transformed normalized read counts between TFX-treated and control samples were calculated, and genes showing more than a twofold enrichment in transposon insertions were included in the summary table (**Fig. 5a**, **Supplementary Data 4**). The Integrative Genomics Viewer (IGV) tool^47^ was used for visualization of the Tn-seq sequencing data.

### NMR analysis

Purified ^15^N-labeled TFX was dissolved in 200 µL of 100 mM phosphate buffer pH 5.8, with 20 µl D_2_O and Trimethyl Silyl Propionate (TSP; 1 mM) as reference, and was introduced in a standard 3-mm NMR tube. All spectra were recorded at 293 K or 280 K on an 800-MHz NEO Bruker spectrometer equipped with a QCP cryogenic probe head. ^1^H, ^15^N HSQC spectra were recorded with 2k x 128 complex points in the ^1^H and ^15^N dimension, for a spectral width of 15.6 x 60 ppm centered at 4.7 and 60ppm, respectively. Decoupling was obtained with a garp4 decoupling sequence during proton acquisition, or was completely omitted. Proton planes of the ^1^H, ^15^N HSQC-TOCSY and HSQC-NOESY spectra were recorded with 2k x 384 points for a spectral width of 12 ppm in both dimensions, with 8 and 32 scans per increment, respectively. Scalar coupling transfer was accomplished during a 60-ms DIPSI2 mixing time, whereas a 200-ms spin lock was used for the dipolar magnetization transfer. The relaxation delay was set to 1s for both spectra. Spectra were repeated at the lower temperature of 280 K in order to shift the water resonance.

Natural abundance ^1^H, ^13^C HSQC, HSQC-DIPSI and HMBC spectra were recorded with standard Bruker pulse programs (hsqcetgpsisp.2, hsqcdietgpsisp.2 and hmbcedetgpl3nd), with 4k x 256 points for a spectral window of 12 x 80 ppm and the ^13^C carrier set at 40 ppm for both HSQC and HSQC-DIPSI experiments, and with 4k x 256 points for a spectral window of 12 x 220 ppm and the ^13^C window centered at 100 ppm for the HMBC experiment. The sample was then lyophilized and resuspended in D2O, and we again recorded the ^1^H, ^13^C HSQC and HMBC spectra at 280K.

All spectra were transformed and analyzed with the Topspin4.1 software.

## DATA AVAILABILITY

Tn-seq sequencing data were deposited in the SRA (BioProject accession n◦ PRJNA1356514). The NMR data for Trifolitoxin were deposited in the Natural Products Magnetic Resonance Database (NP-MRD; www.np-mrd.org) and can be found at NP0351953. All other data generated or analyzed during this study are included in this published article and its supplementary information files.

## AUTHOR CONTRIBUTIONS

D.Y.T. and S.D. designed the study. D.B. performed genome mining and analysis of tfx-like BGCs with input from D.Y.T. and S.D.; D.Y.T., C.P., F.W, K.G., A.L., and S.D. performed biochemical and microbiological experiments. A.L., M.S., and S.Z. performed MS analyses. G.L. performed NMR experiments. D.N., T.T., and P.M. obtained Tn-seq data. All authors analyzed and interpreted the data. D.Y.T. and S.D. created the figures and wrote the manuscript with input from S.Z., G.L., and P.M.

## ACKNOWLEDGMENTS

We are grateful to Prof. Alexander Mankin (University of Illinois at Chicago, USA), Prof. Nora Vazquez-Laslop (University of Illinois at Chicago, USA), Prof. Yury Polikanov (University of Illinois at Chicago, USA), and Prof. Dmitry Ghilarov (Oriel College, University of Oxford, UK) for useful insights. We thank Prof. Matthew Henke and Ms. Ngoc Pham (University of Illinois at Chicago, USA) for the collection of HR-MS data for processed TFX, and Dr. Aleksei Kulikovsky for the help with the large-scale purification of TFX. We thank Ms. Elena Aleksandrova (Polikanov Lab, UIC) for providing purified deacylated tRNA^Arg^. We thank MetaboHub-MetaToul (Metabolomics and Fluxomics facilities, Toulouse, France), part of the French National Infrastructure for Metabolomics and Fluxomics for access to the NMR facility and the Moscow State University Development Programme APG 5.13 for access to the mass spectrometry facility. The authors thank Prof. Victor Zgoda and the Core Facilities at Institute of Biomedical Chemistry, Russian Academy of Sciences for assistance with MS data acquisition for nucleoside samples, and the I2BC sequencing platform (CNRS, Gif-sur-Yvette, France) for generating sequencing data.

This work was supported by the INRAE EXPLOR’AE program, a part of the France 2030 initiative, grant ANR-24-RRII-0003 to S.D. and G.L. and Illinois State startup funds to D.Y.T.

---

**Extended Data Figure 1.**
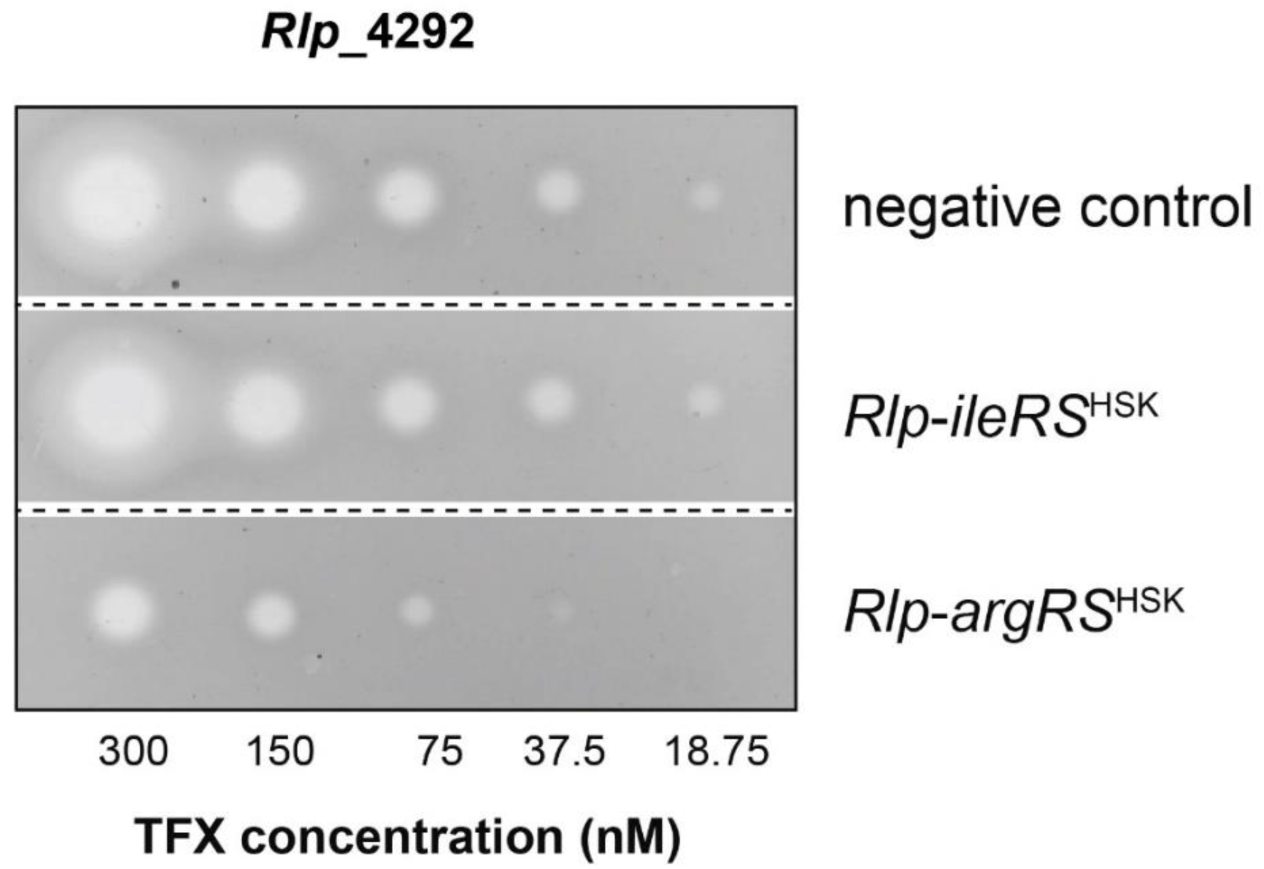
Inhibition zones from 5 μL aliquots of serial two-fold TFX dilutions spotted onto lawns of *Rlp_*4292 derivatives grown on medium containing 1 mM IPTG. Overexpression of *Rlp*-*argRS*^HSK^ but not *Rlp*-*ileRS*^HSK^ makes bacteria less sensitive to TFX.

**Extended Data Figure 2.**
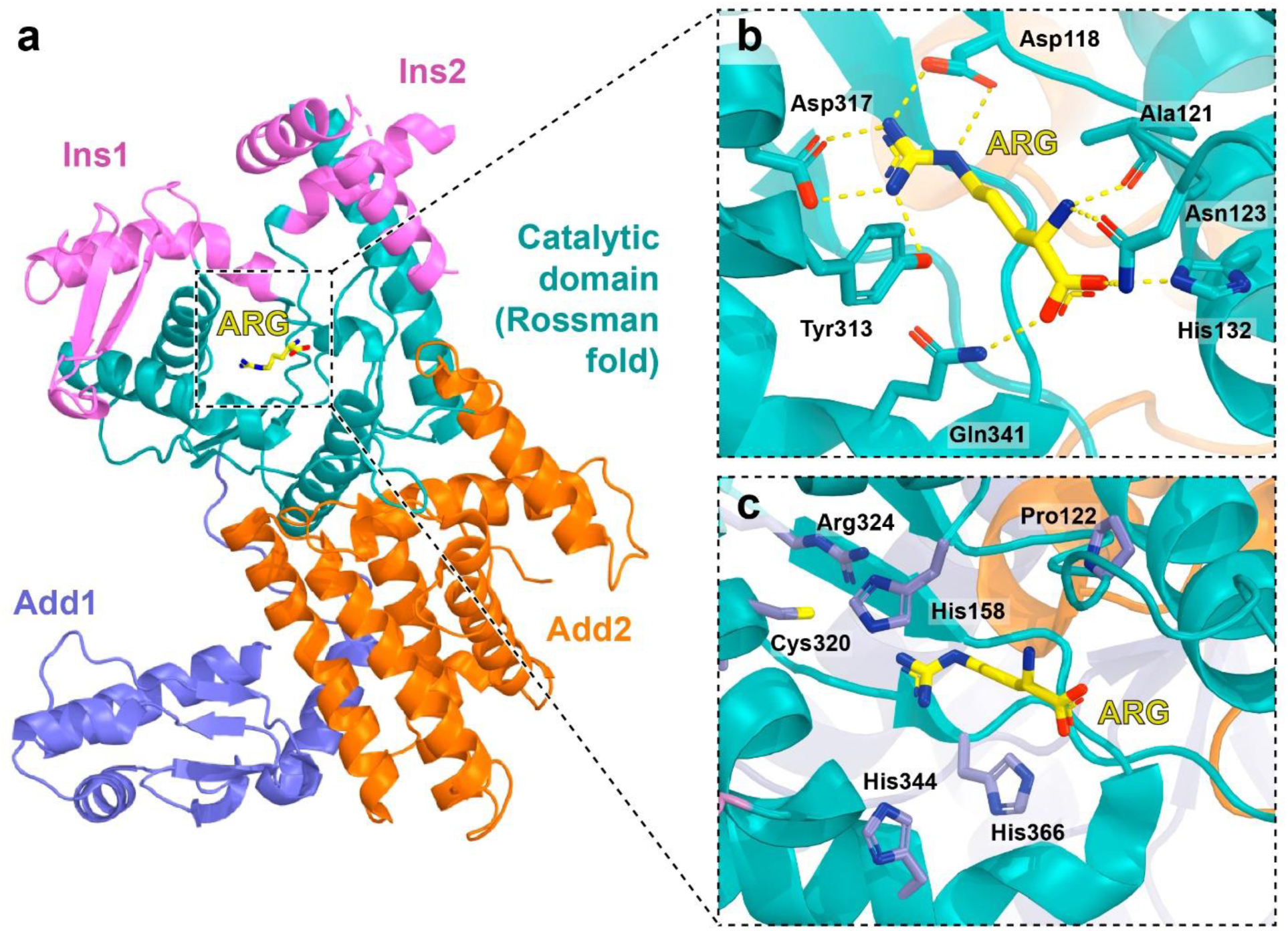
a. Structure of *Eco*-ArgRS^HSK^ in complex with arginine. (ARG) (PDB ID: 4OBY^18^). The domains are colored according to ref. ^49^: Add1 – ArgRS-specific additional N-terminal domain involved in tRNA recognition, Add2 – additional domain 2, Ins1 and Ins2 – insertion domains. b. Residues involved in ARG coordination *via* hydrogen bonds. **c.** Residues forming the ARG-binding site but not involved in the ligand coordination.

**Extended Data Figure 3.**
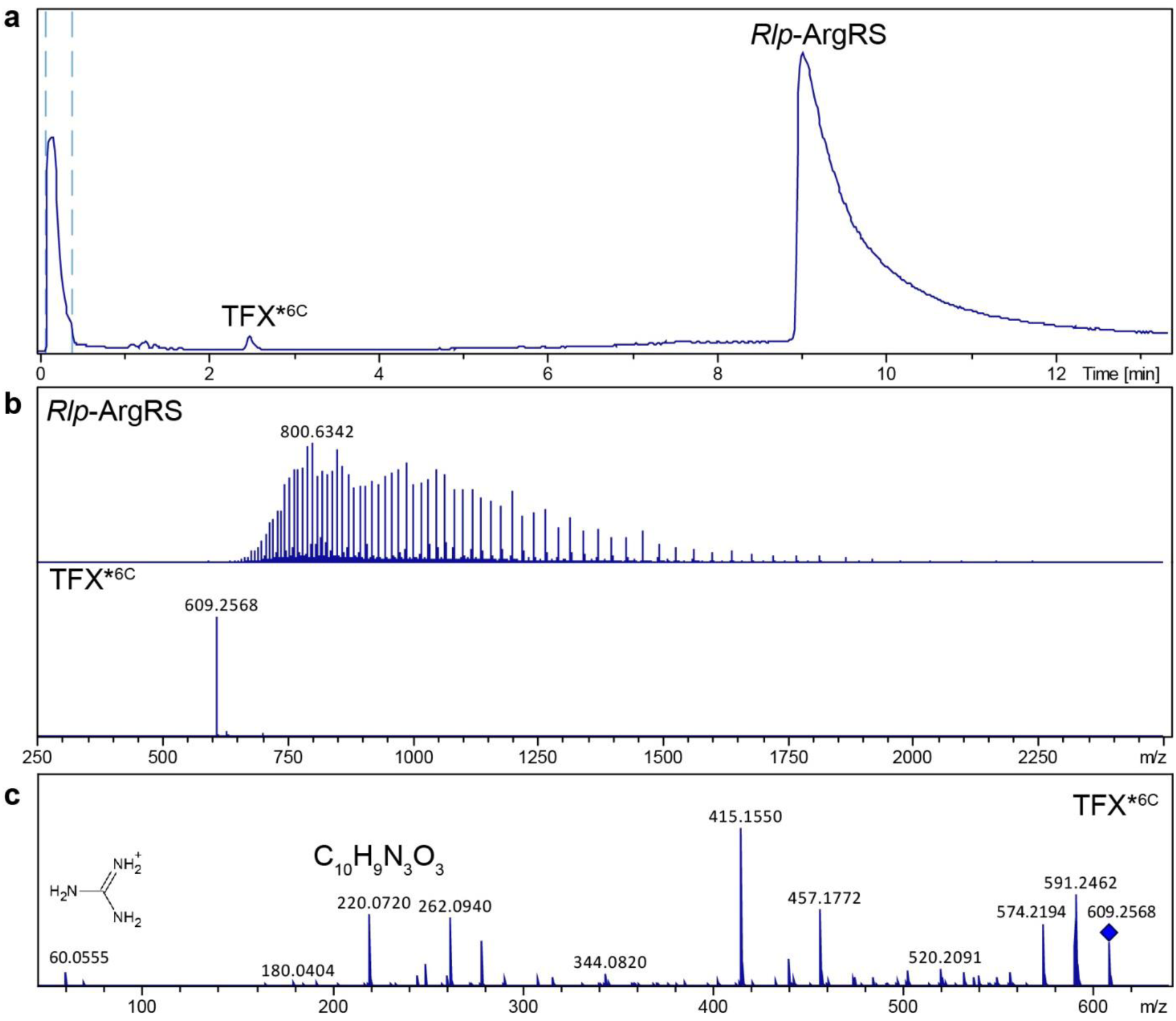
Native LC-MS analysis of Rlp-ArgRS purified from TFX-treated cells under native conditions. **a.** Total ion chromatogram showing the distinct peaks corresponding to TFX*^6C^ (RT 2.5 min) and *Rlp*-ArgRS^HSK^ (RT 9 min). **b.** MS spectra of TFX^*6C^ and *Rlp*-ArgRS^HSK^. **c.** MS/MS spectrum of TFX^*6C^: [M+H]^+^ at *m/z* 609.26.

**Extended Data Figure 4.**
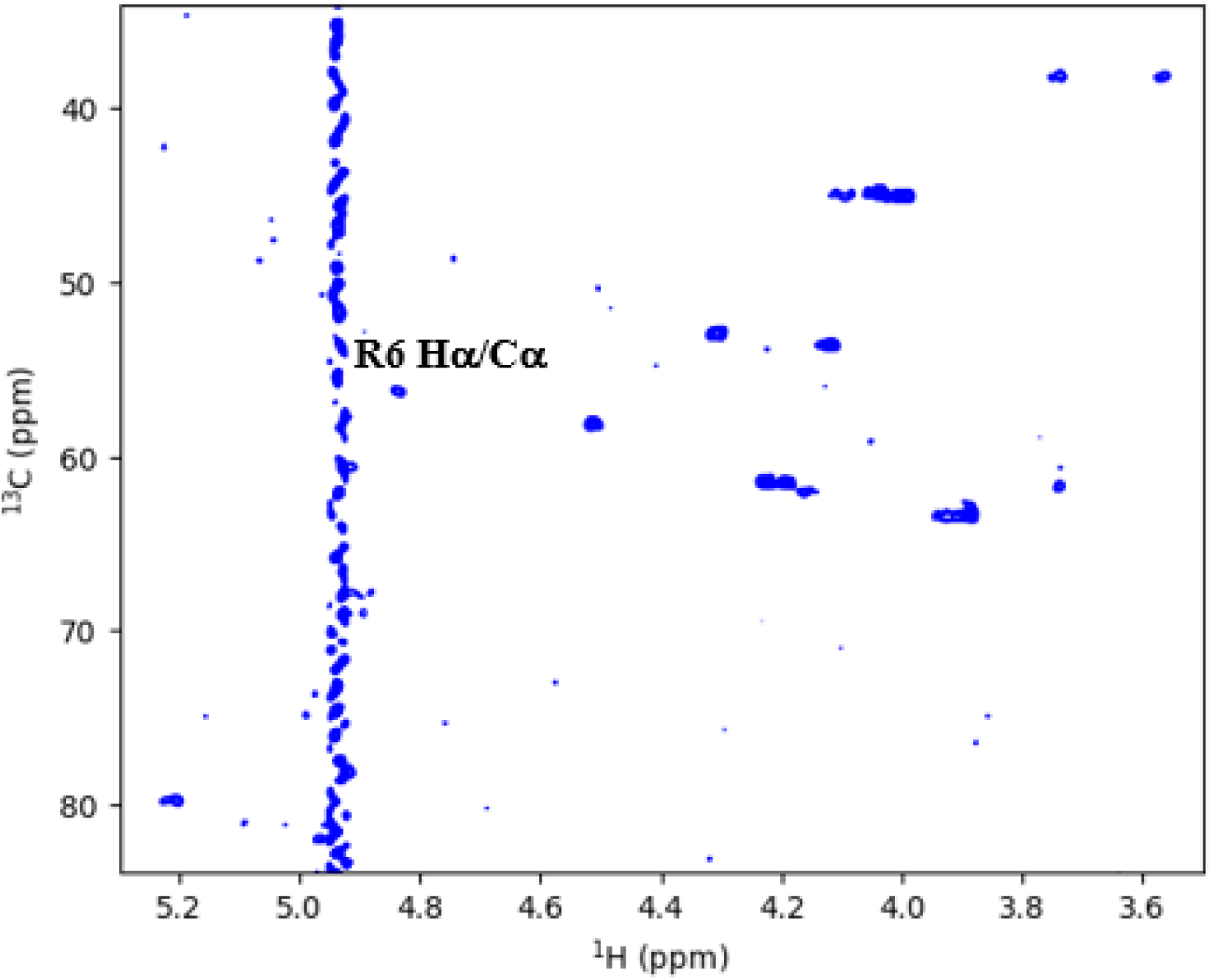
^1^H, ^13^C HSQC spectrum of TFX in D_2_O at 280. **K.** The Hα resonance of Arg6 at 4.83ppm can be identified as the cross peak at a ^13^Cα value of 56.2ppm.

**Extended Data Figure 5.**
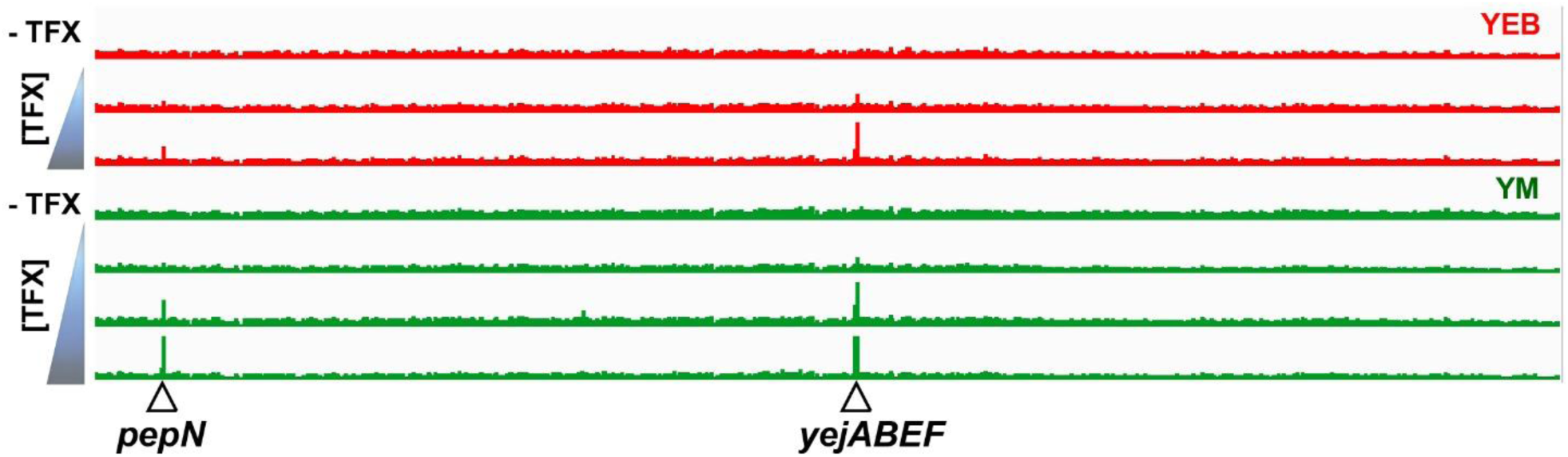
IGV view of Tn-seq sequencing data upon TFX treatment in two different cultivation media (YEB and YM) for the whole genome of *Rlp*_4292. The histograms indicate the abundance of mutants in the Tn-seq population for the indicated conditions. The arrows indicate the position of the *pepN* gene and *yejABEF* operon, highlighting a dose-dependent increase of Tn insertion frequencies with rising TFX concentrations.

**Extended Data Figure 6.**
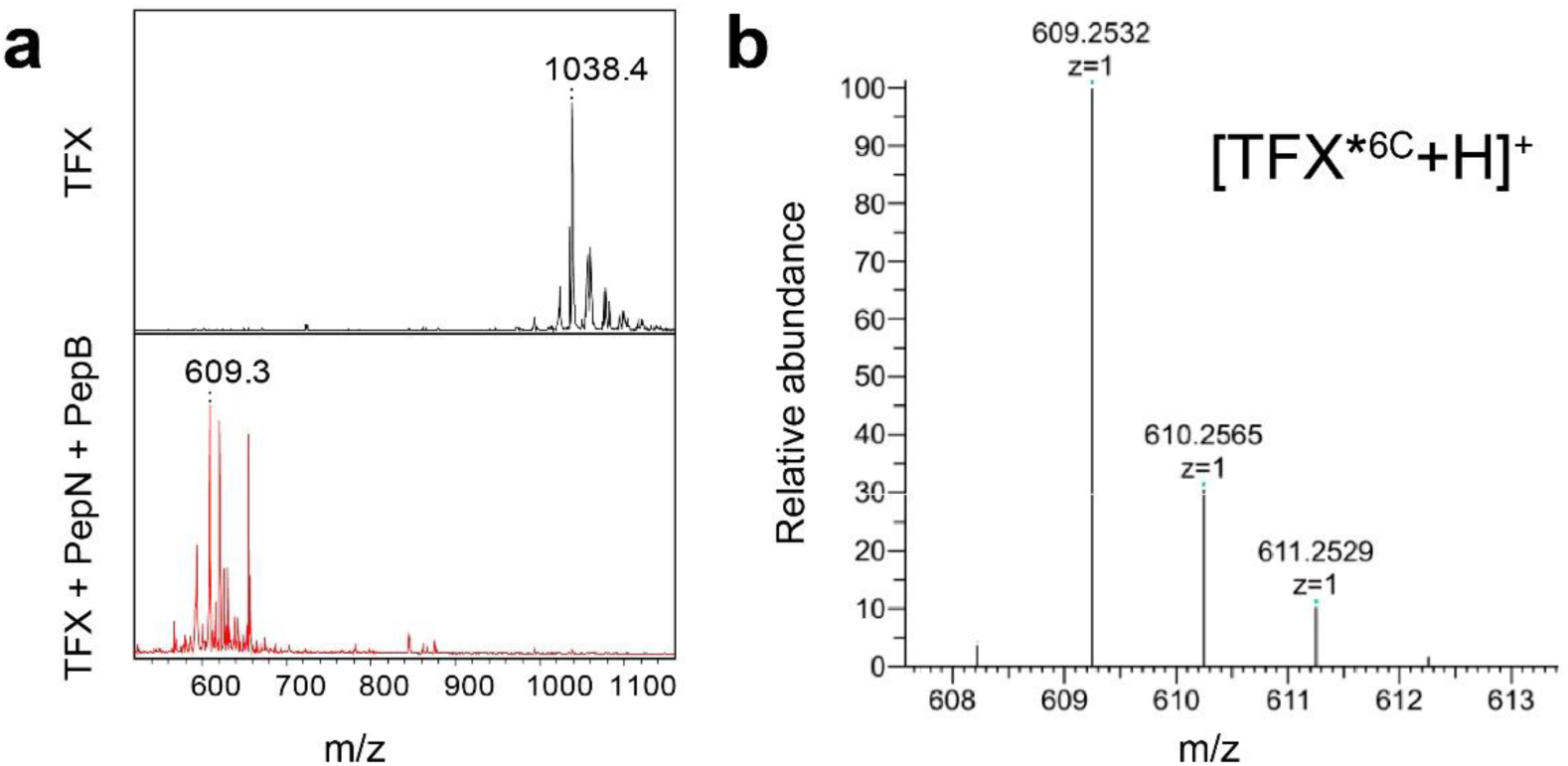
MS spectra of proteolytically processed TFX. **a.** MALDI-TOF MS spectra of mature TFX and the products of its *in vitro* digestion by purified *E. coli* aminopeptidases PepB and PepN. Note the appearance of the [M+H]^+^ ion at *m/z* 609.3 corresponding to the N-terminally processed TFX. **b.** High resolution ESI-MS spectrum of proteolytically processed TFX (TFX*^6C^). The *m/z* values for each mass ions are indicated. Theoretical MW: 608.2489; Experimental MW: 608.2460; Mass error (ppm): - 4.77.

**Extended Data Fig. 7.**
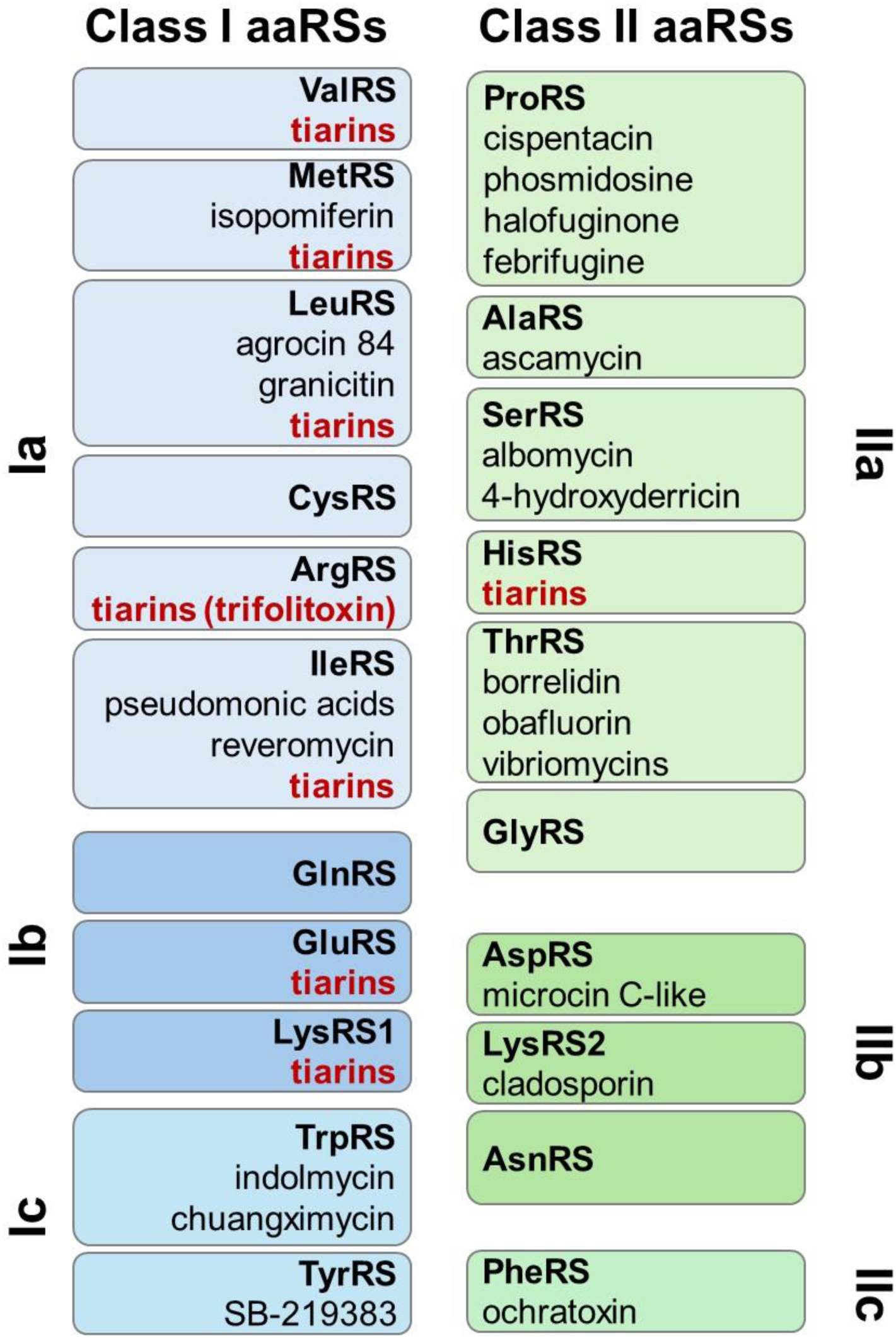
aaRSs targeted by tiarins and other known natural compounds.

**Extended Data Table 1.**
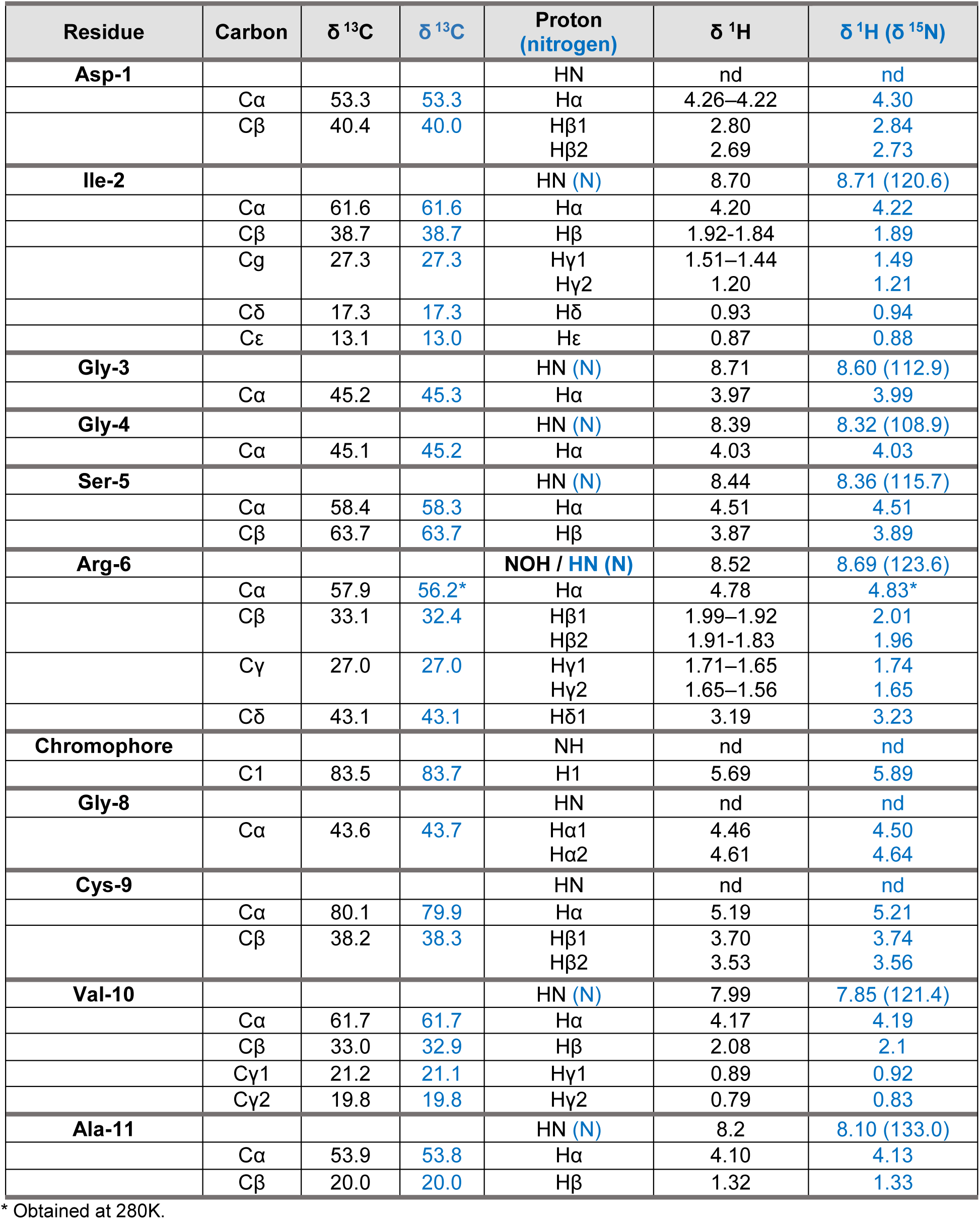
NMR chemical shifts reported by Lethbridge *et al.*^8^ and in the present study.

## Notes

### Competing Interest Statement

The authors have declared no competing interest.

