## Supplementary Materials for "Tiarins, a diverse family of natural Trojan-horse aminoacyl-tRNA synthetase inhibitors discovered by genome mining"

for

**Supplementary Discussion**

The $^{1}H$, $^{15}N$ HSQC spectrum of the $^{15}N$-labeled TFX sample shows 7 cross peaks with $^{15}N$ chemical shifts in the [110-130ppm] range (**Supplementary Fig. 7**), corresponding to amide moieties. No signal of an $NH_2$ group can be seen, but we do observe a cross peak at 83.6ppm corresponding to an arginine side chain resonance. Assignment of the backbone resonances was done based on the proton planes of a TOCSY- and NOESY-HSQC spectrum, and resulted in the assignment of the Ile2-Arg6 and Val10-Ala11 unmodified amino acids of TFX (**Supplementary Fig. 7**). To further ascertain that all amide nitrogens were directly protonated (and not hydroxylated, as proposed for R6 in the original structure of TFX), we recorded the $^{1}H$, $^{15}N$ HSQC spectrum without $^{15}N$ decoupling in the acquisition. The $^{1}J$ coupling constant of 90Hz found for all 7 residues (Arg6 included) confirms the absence of modification for these amide moieties (**Fig 4d**). The Hα proton of Arg6 coincides with the water resonance at 4.8ppm, but spectra at 280K confirmed its presence (**Supplementary Fig. 8**). When we lyophilized the sample and resuspended it in $D_2O$ at this lower temperature, the Hα/Cα correlation of the Arg6 remained visible at 4.83/56.2ppm (**Extended Data Fig. 4**).

The $^{1}H$, $^{13}C$ HSQC and TOCSY-HSQC spectra (at natural abundance) further showed the N-terminal Asp residue and several remaining $CH_2$ groups (**Supplementary Fig. 9**). In agreement with the assignment of Lethbridge *et al*., one of the latter with its $^{13}C$ frequency of 43.7ppm could be assigned to the modified Gly8, and a second one at 38.2ppm to the cyclized Cys9 residue. After lyophilization and resuspension in $D_2O$, we found two further resonances downfield of the residual water line at 4.8ppm. One cross peak at 5.22/79.9ppm connected with the $CH_2$ signal at 38.2ppm, and could be assigned to the cyclized Cys in position 9 (**Supplementary Fig. 10**). The second singlet resonance, at $d^{1}H$ 5.89/$d^{13}C$ 83.7ppm, was assigned by Lethbridge *et al*. to the vinyl proton of the chromophore coming from the modified glutamine.

In conclusion, whereas our assignments are generally in good agreement with those of Lethbridge et *al*. (**Extended Data Table 1**), we do not find any evidence for the proposed modification of Arg6. Hence, as agreement with the mass spectrometry data is no longer guaranteed when one eliminates the hydroxyl group on the Arg6 amide nitrogen, further structural studies will be needed to establish the molecular formula of TFX.

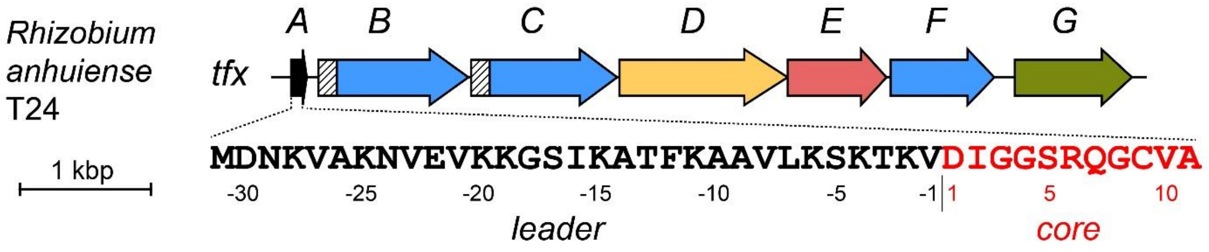


| Gene | Protein MW (kDa) | Protein pI | Domain prediction (Interpro) | Proposed function |
|---|---|---|---|---|
| *tfxA* | 4.41 | 9.24 | - | Precursor peptide |
| *tfxB* | 42.50 | 5.19 | SagB-type dehydrogenase domain (IPR020051) | PTM (chromophore installation?) |
| *tfxC* | 39.86 | 5.01 | Nitroreductase-like (IPR000415) | PTM (chromophore installation?) |
| *tfxD* | 44.87 | 9.46 | Transmembrane protein (no domains found) | Export of mature product |
| *tfxE* | 28.04 | 9.25 | YcaO-like domain (IPR003776) | PTM (thiazoline installation?) |
| *tfxF* | 29.49 | 4.88 | Nitroreductase-like (IPR000415) | PTM (chromophore auxillary modifications?) |
| *tfxG* | 28.82 | 5.30 | Aminoglycoside phosphotransferase (IPR002575) | Production of trifolitoxin isomers? |

**Supplementary Figure 1 |** Characteristics of proteins encoded in the *tfx* biosynthetic gene cluster.

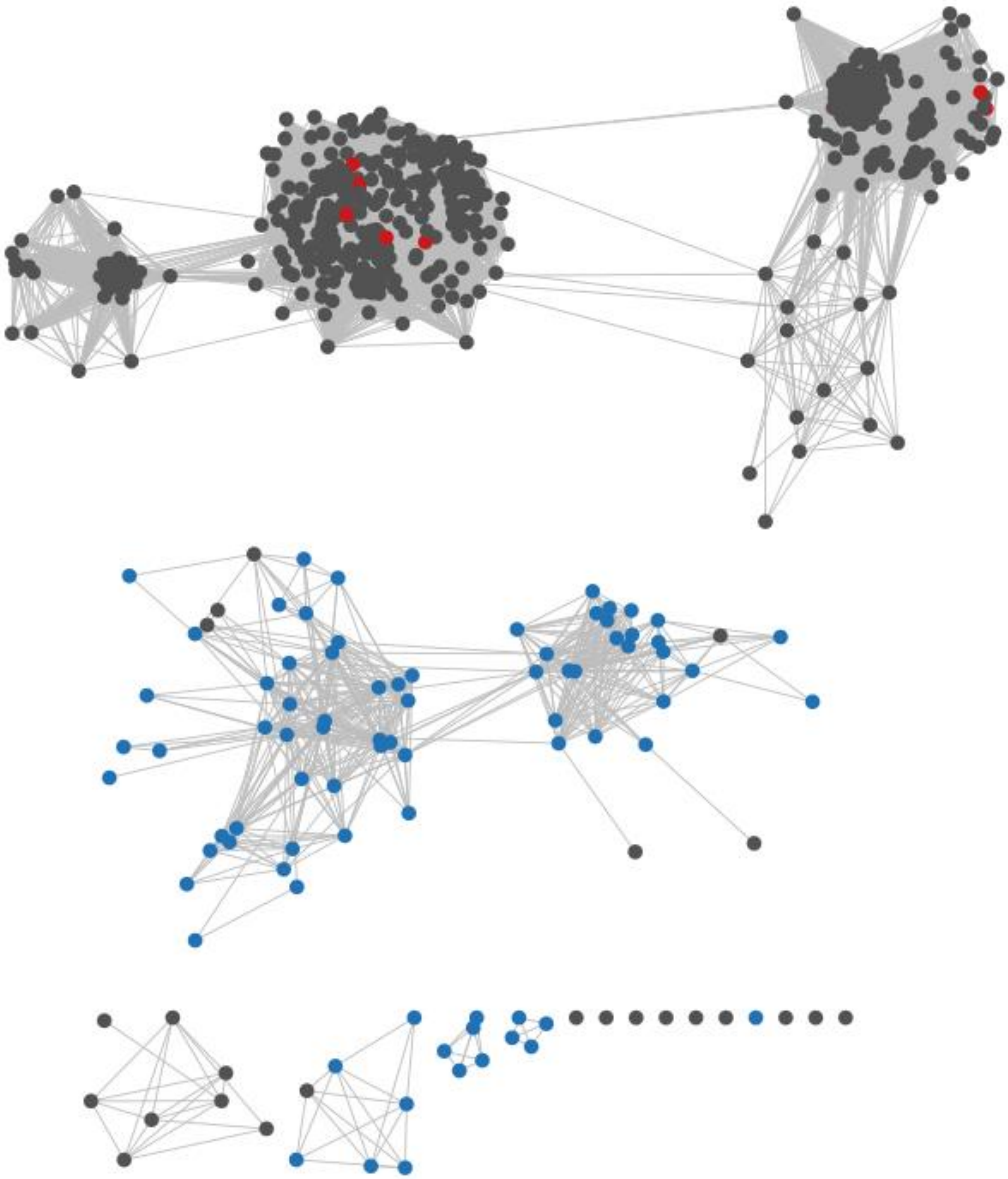

**Supplementary Figure 2. Sequence similarity network of representative TfxC homologs**. Edges with an alignment score (negative log of e-value) below 32 were removed. Colors indicate the genomic distance between genes encoding TfxC and TfxB homologs: grey, TfxB not detected; blue, distance ≤ 2.7 kb; red, distance between 2.4 and 6.5 Mb. The final dataset for phylogenetic tree construction included only TfxB–TfxC homolog pairs whose genes are separated by ≤2.7 kb.

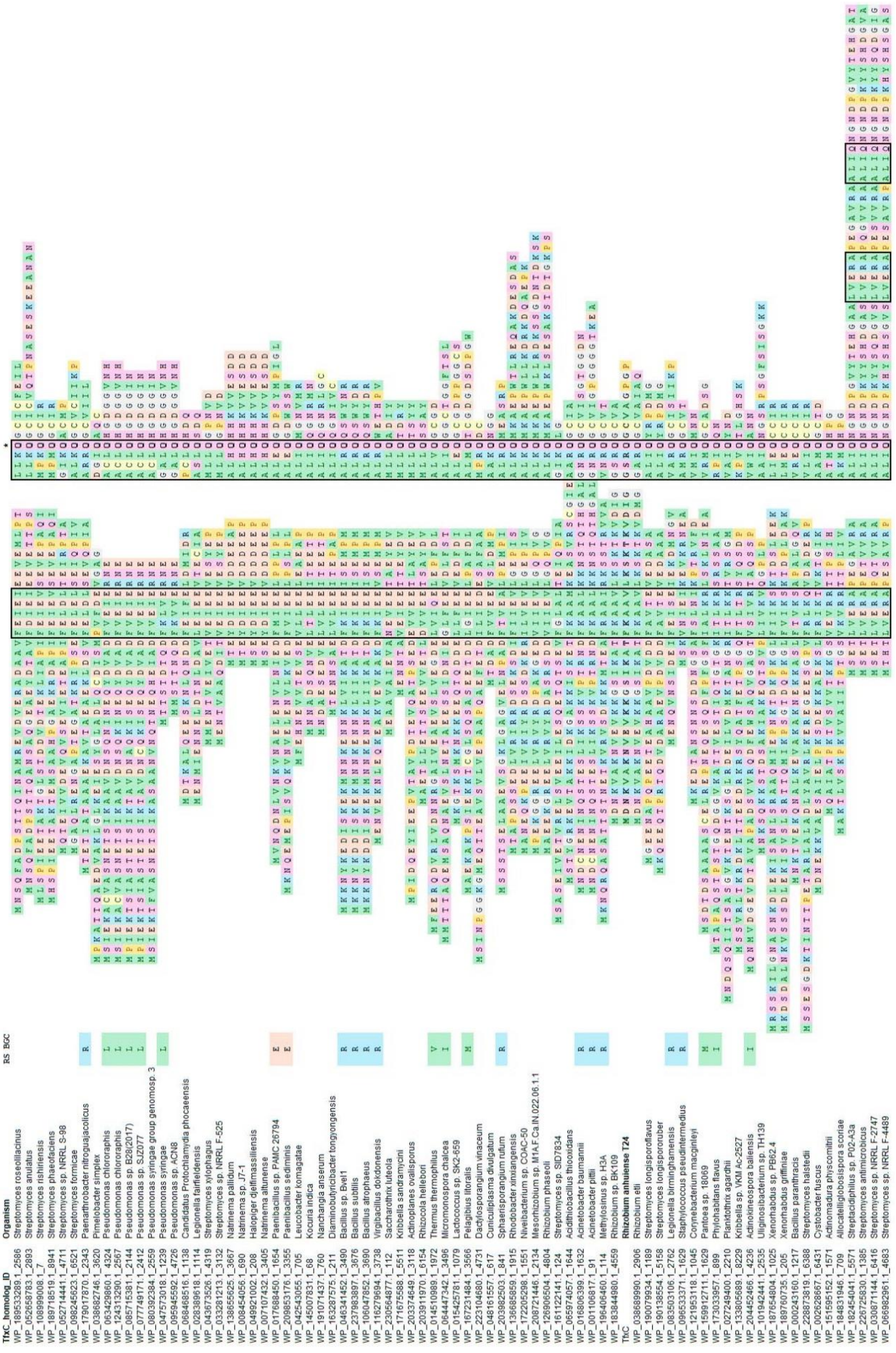

**Supplementary Figure 3 |** Multiple alignment of the amino acid sequences of the precursor peptides encoded in *tfx*-like BGCs. The color scheme reflects the chemical properties of the amino acid side chain. Motifs 1 and 2 are shown in black squares. An asterisk indicates the conserved Gln residue in motif 2. Note that four tiarin BGCs from *Streptomyces* shown in the lower part of the alignment encode cassette precursor peptides with two leader and two core parts fused into a single open reading frame.

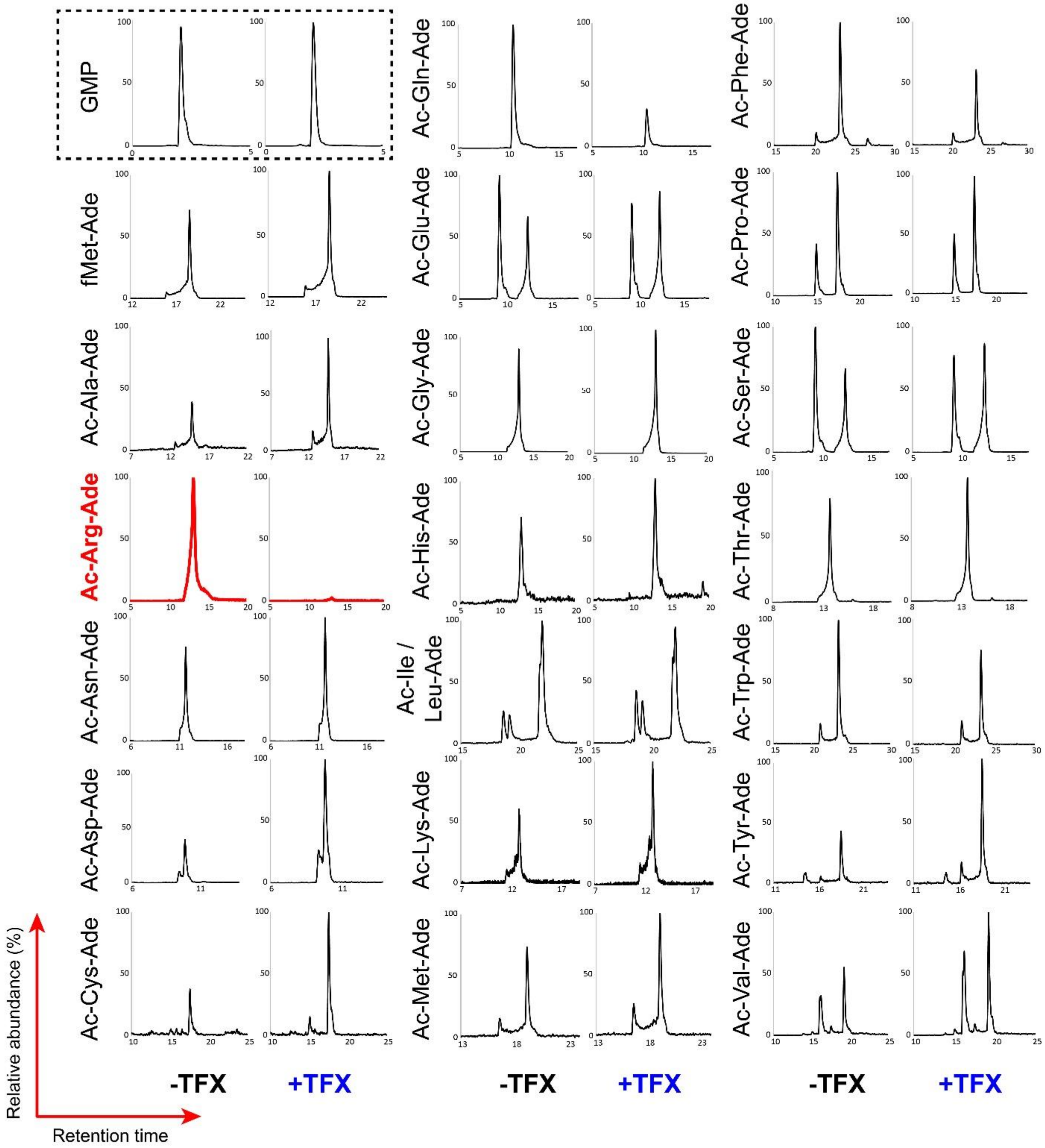


**Supplementary Figure 4 |** LC-MS analysis of RNase-digested acetyl-aminoacyl-tRNAs fragments. Extracted ion chromatograms (EICs) for the protonated Ac-aminoacyl-A76 are shown for one out of three experimental replicates in two conditions (no TFX, +TFX). Signals from $^{13}C$-GMP (shown in a dashed frame) were used for normalization within the pair of samples.

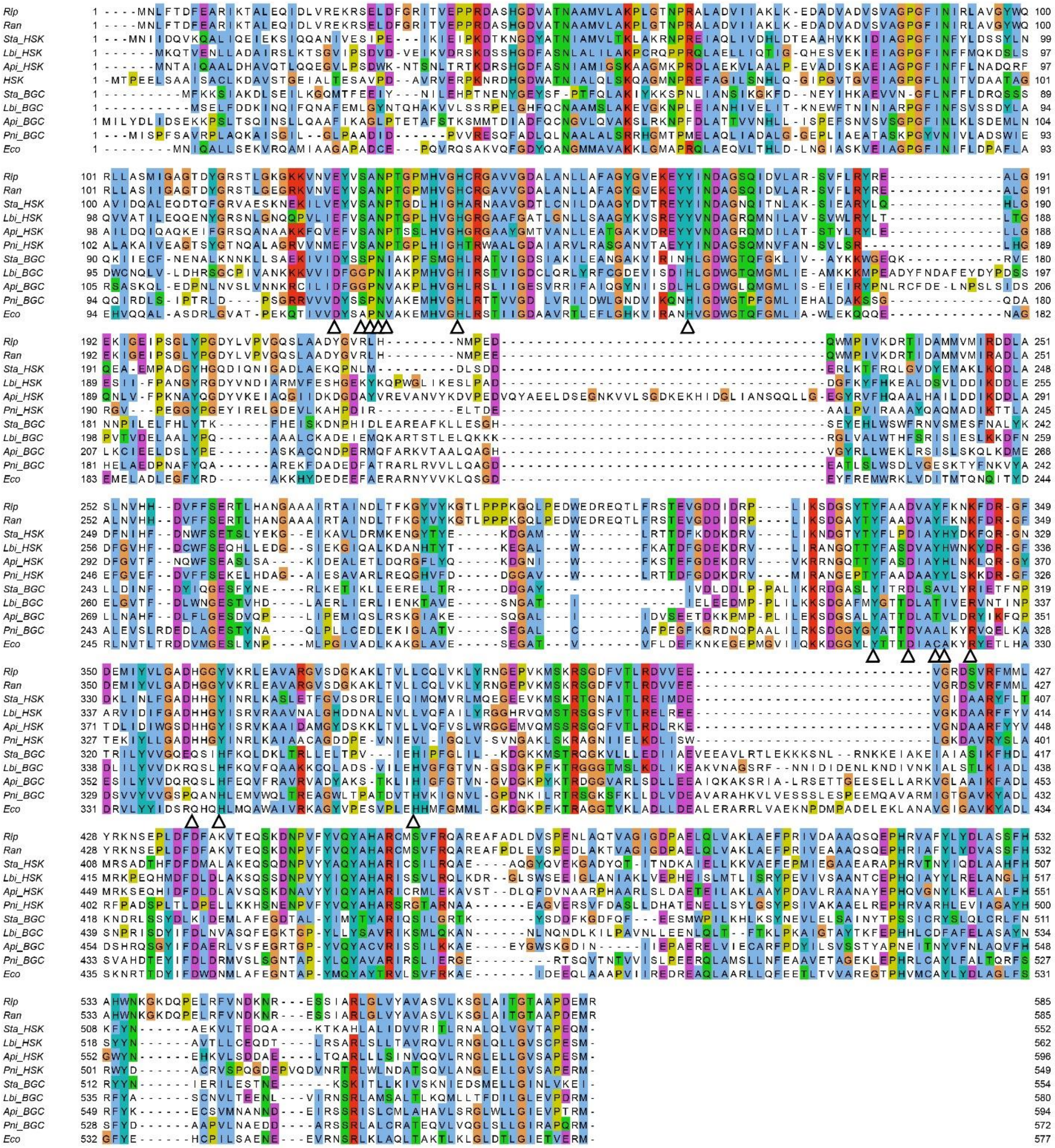


**Supplementary Figure 5 |** Multiple sequence alignment of ArgRSs from the genomes of *Rhizobium leguminosarum* 4292 (Rlp), *Rhizobium anhuiense* T24 (Ran), *Staphylococcus* sp. MI 10-1553 (Sta), *Legionella birminghamensis* DSM 19232 (Lbi), *Acinetobacter pittii* DSM 25618 (Api), *Paenarthrobacter nitroguajacolicus* xvA3 (Pni), and *Escherichia coli* BW25113 (Eco) built by the CLUSTAL W algorithm[1]. For the strains encoding both the “housekeeping” and BGC-associated ArgRSs, those are labeled as HSK and BGC, respectively. The alignment is colored using the Clustal scheme, reflecting the chemical properties of the amino acid side chains. The alignment was visualized using Jalview.[2] White arrowheads indicate the residues shown in Fig. 3b.

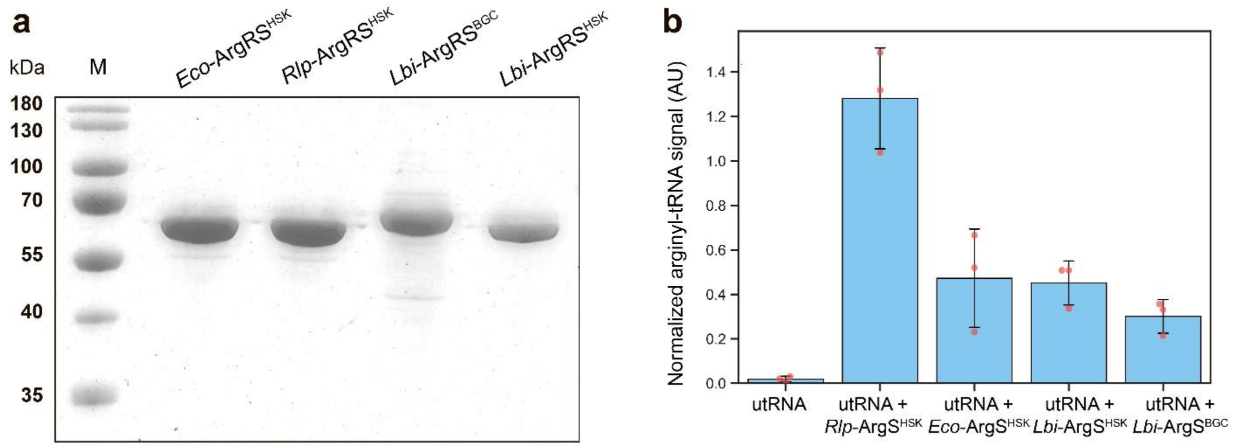


**Supplementary Figure 6 | a.** SDS-PAGE analysis of heterologously expressed and affinity-purified 6His-tagged variants of ArgRSs of different origin. M – protein molecular weight marker. **b***.* *In vitro* aminoacylation of uncharged tRNAs (utRNA) of *Rlp*_4292 by four heterologously produced ArgRSs of different origin. The standard deviation for three independent replicates is shown.

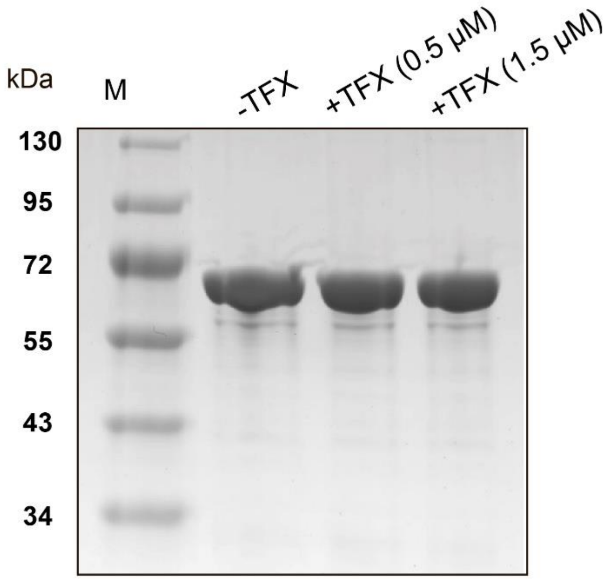


**Supplementary Figure 7 |** SDS-PAGE analysis of *Rlp*-ArgRS$^{HSK}$ samples affinity purified from the control *Rlp*_4292 culture and the cultures treated with TFX in two different concentrations.

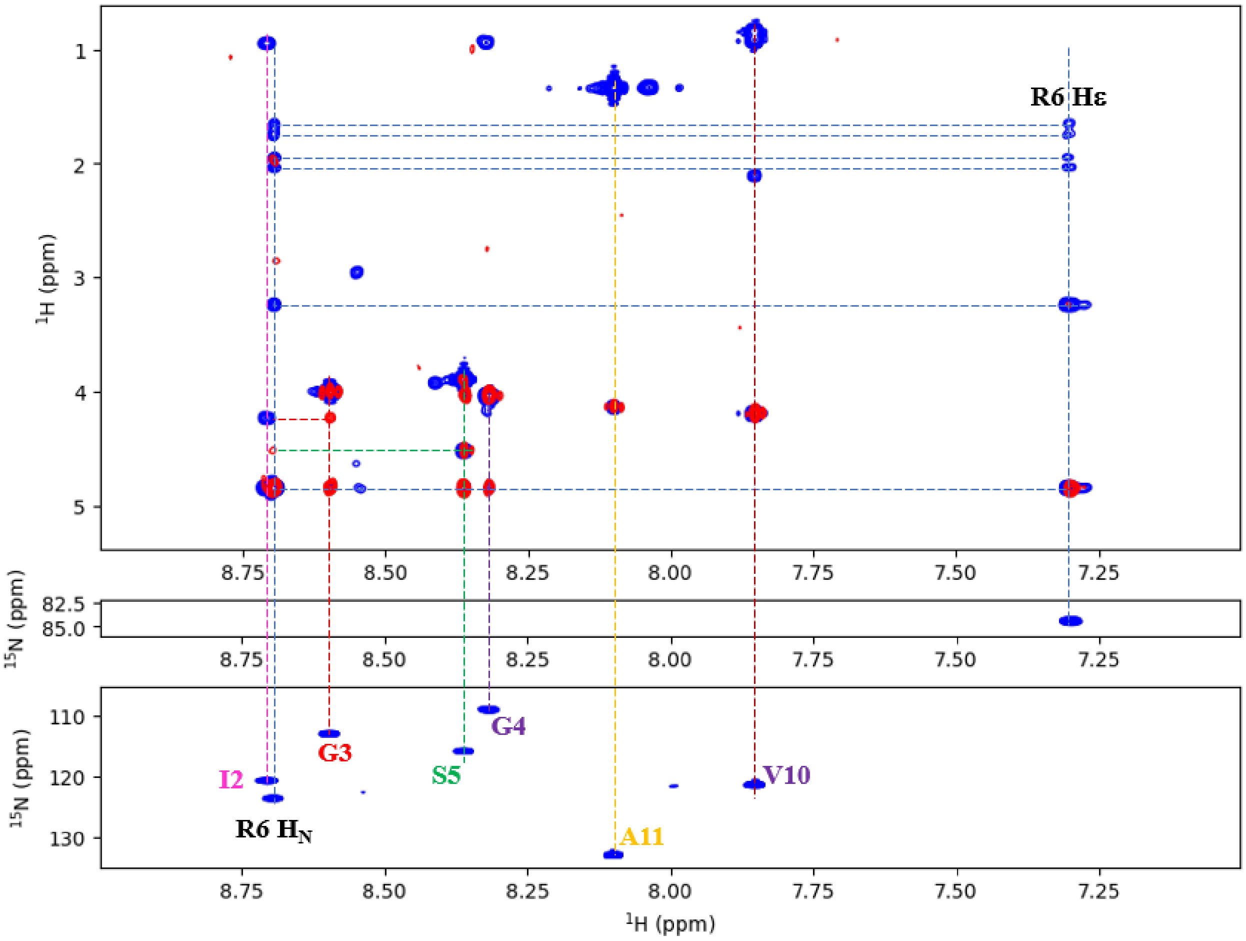


**Supplementary Figure 8 |** Assignment of the peptidic part of TFX. (top) Planes of the (blue) TOCSY- and (red) NOESY-HSQC spectra with the corresponding assignments. (middle) Cross peak of the Arg6 side chain HN group. (bottom) $^{1}H$, $^{15}N$ HSQC spectrum of the backbone region of TFX.

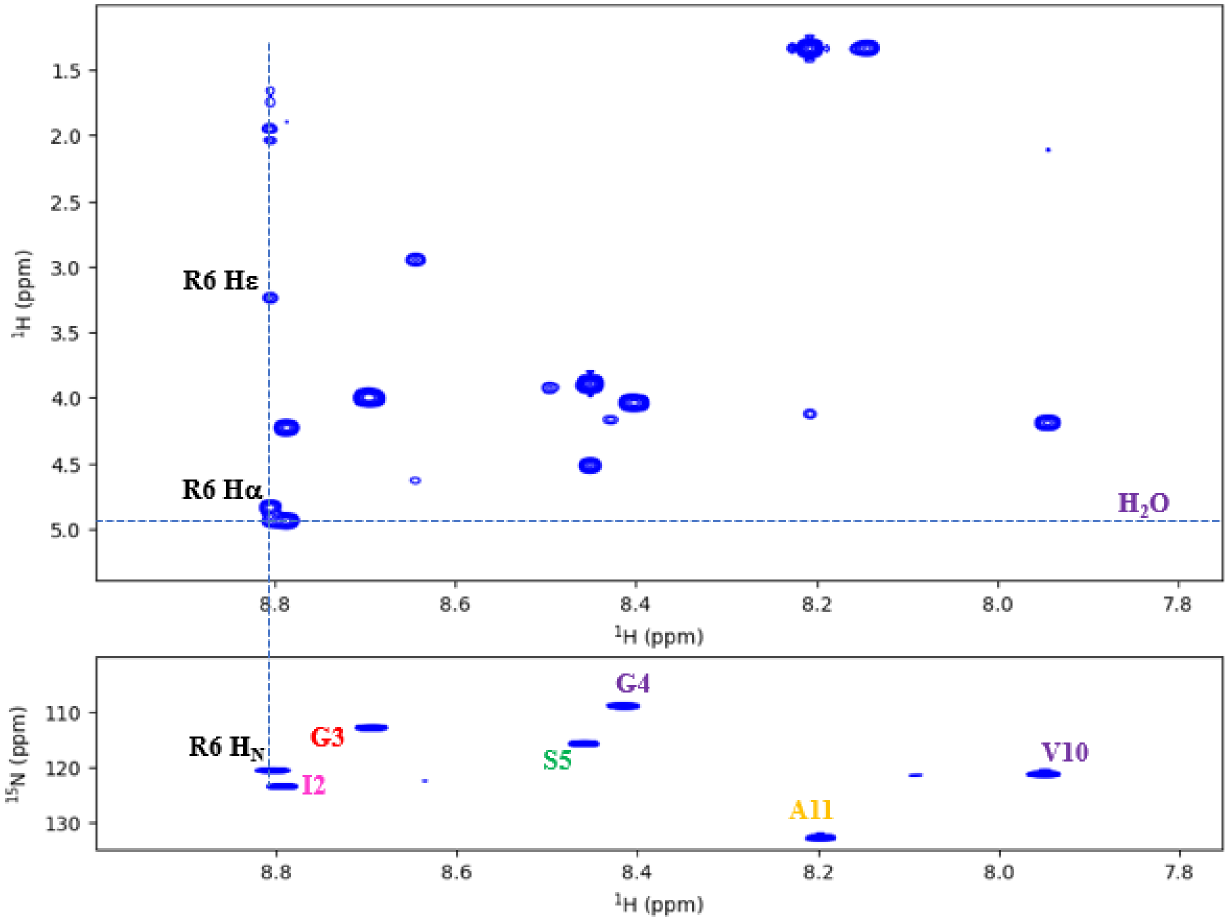


**Supplementary Figure 9 |** Assignment of the peptidic part of TFX at 280K. (top) Proton plane of the TOCSY- HSQC spectrum. The water signal shifts downfield at the lower temperature, allowing the distinction of the R6 Hα proton. (bottom) $^{1}H$, $^{15}N$ HSQC spectrum of the backbone region of TFX with the corresponding assignments.

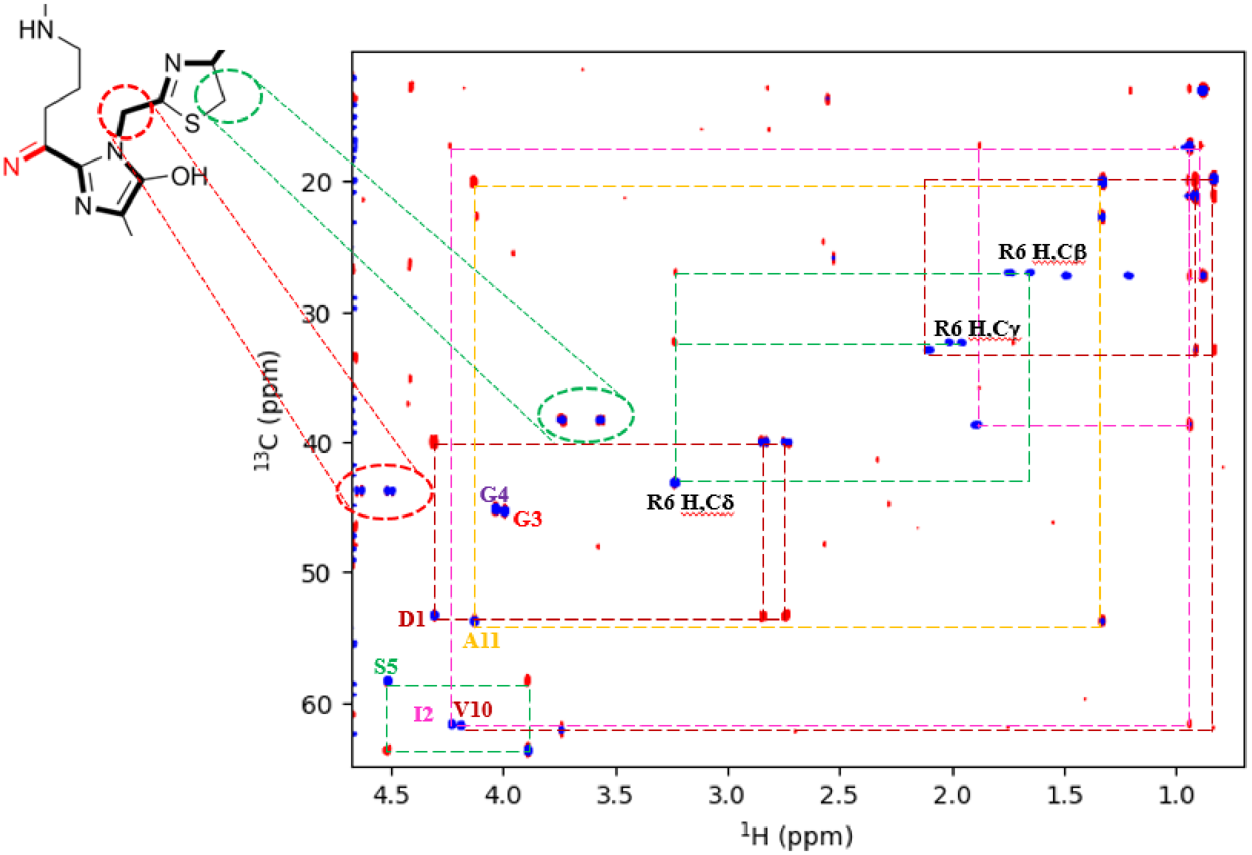


**Supplementary Figure 10 |** $^{1}$H, $^{13}$C HSQC spectrum of TFX at natural abundance. A first $CH_2$ group with $^{13}$C frequency of 43.7ppm (red circle) could be assigned to the modified G8, and a second one at 38.2ppm (green circle) to the cyclized C8 residue. The molecular structure was adapted from Lethbridge *et al.*[1].

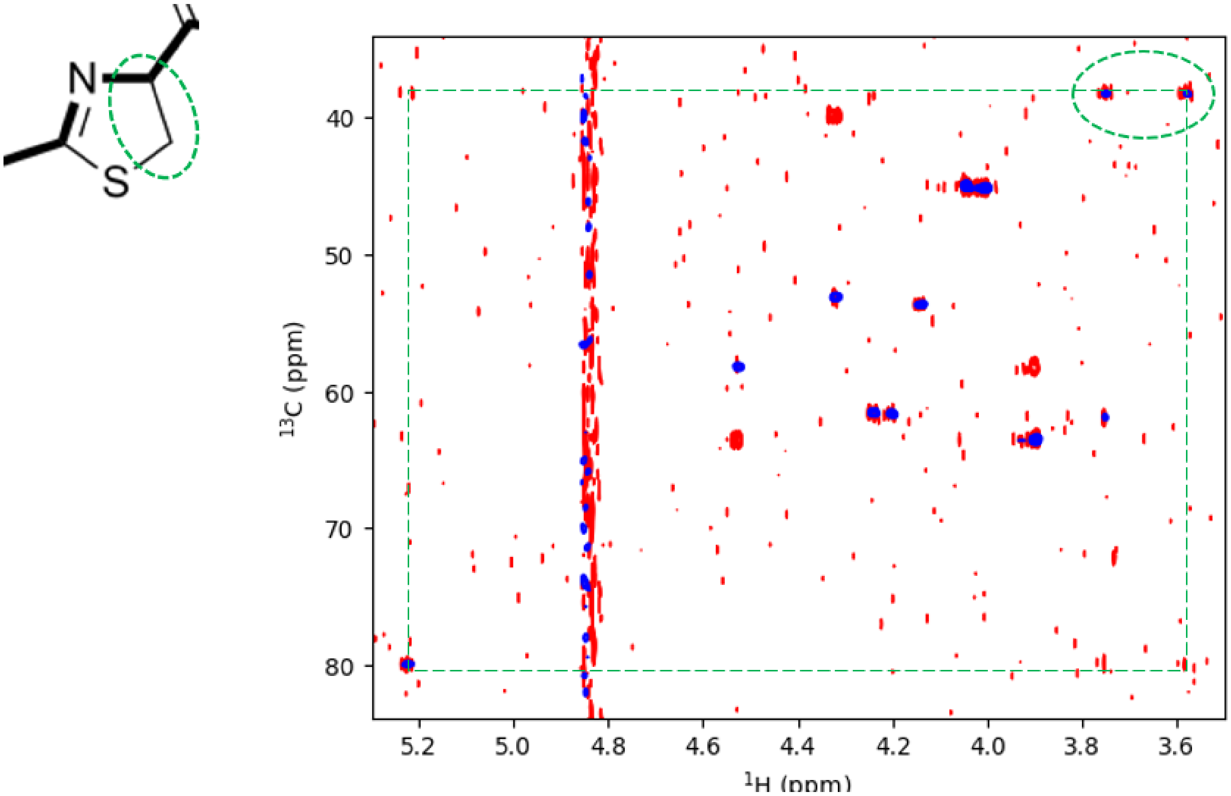


**Supplementary Figure 11 |** $^{1}H$, $^{13}C$ HSQC (blue) and TOCSY-HSQC (red) spectra of TFX in $D_2O$. The $CH_2$ group with $^{13}C$ frequency of 38.2ppm (green circle) connects to the Hα proton of the cyclized C8 residue. The molecular structure was adapted from Lethbridge *et al.*[1].

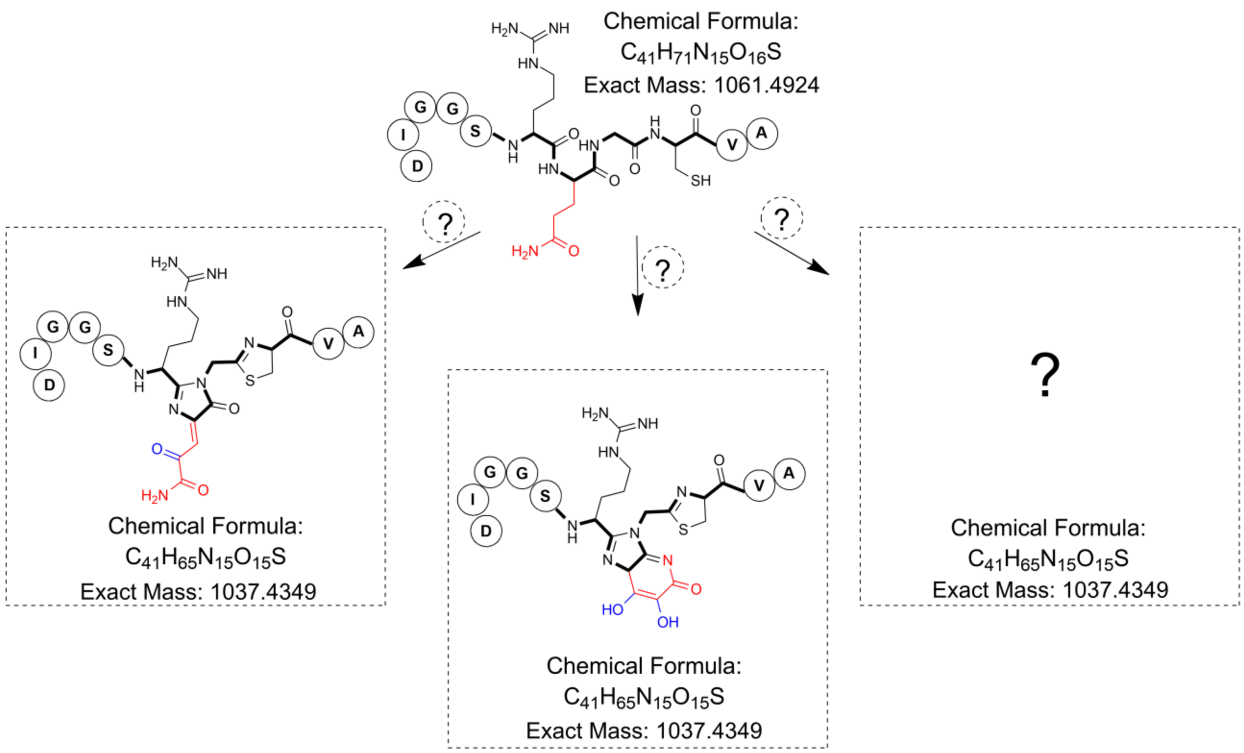


**Supplementary Figure 12 |** Proposed alternative structures of the TFX fluorophore.

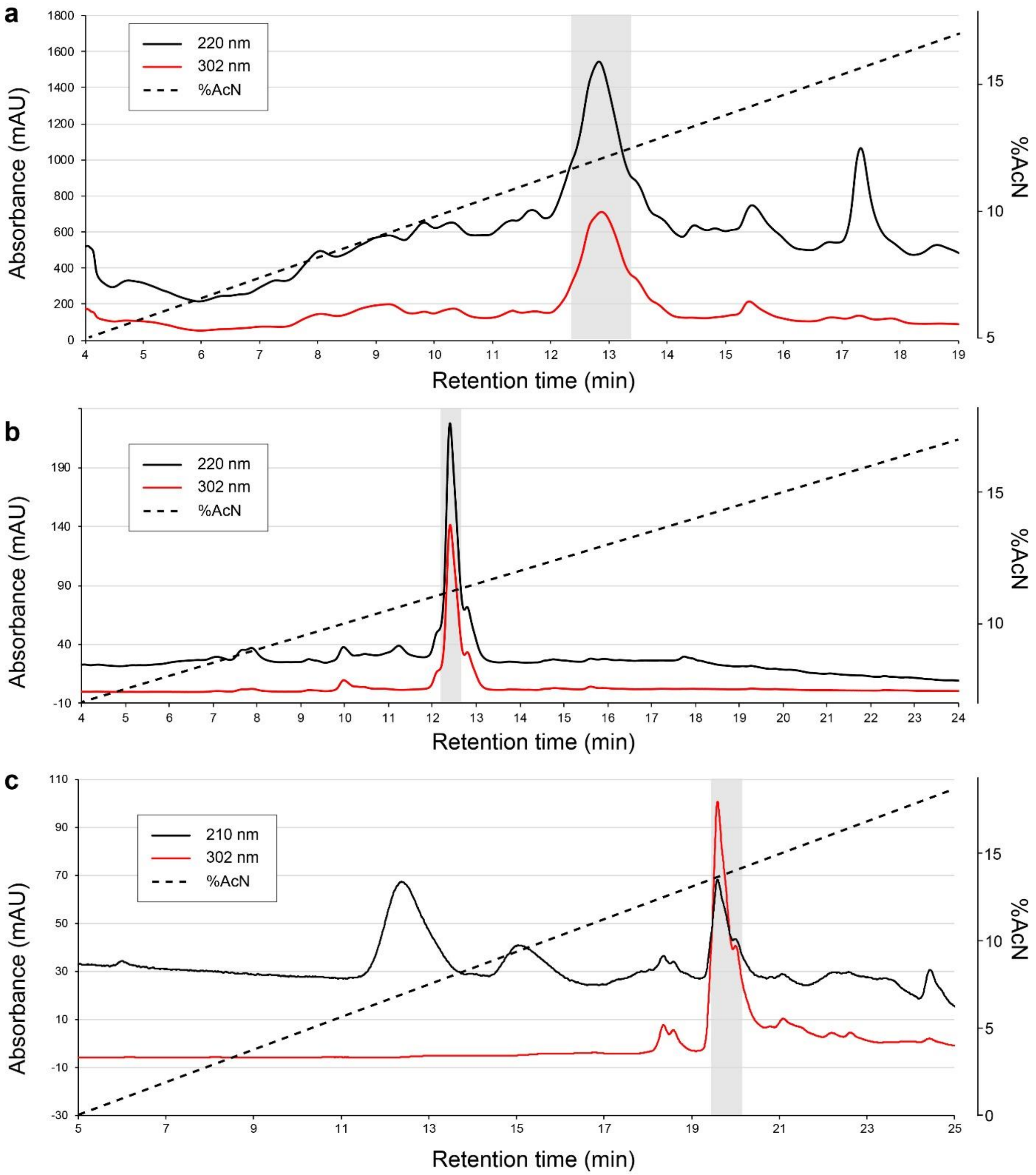


**Supplementary Figure 13 | Purification of TFX and TFX$^{*6C}$ by HPLC a.** HPLC profile of the crude spent medium extract after *R. anhuiense* T24 cultivation. The TFX-containing fraction collected is shown on the gray background. **b.** HPLC profile of the active TFX-containing fraction from **a**. (second round of purification). The TFX-containing fraction collected is shown on the gray background. **c.** HPLC profile of the in vitro reaction of TFX cleavage by aminopeptidases PepN and PepB. The fraction containing TFX$^{*6C}$ is shown on the gray background. **c.** HPLC profile of the *in vitro* reaction of TFX cleavage. Peak containing TFX*6 is indicated

**Supplementary Table 1 | Locus tags of aaRS-encoding genes in the bacterial genomes harboring tiarin BGCs with a putative self-resistance aaRS gene.**

| Assembly accession # | Organism | aaRS specificity | BGC-associated aaRS's locus tag | Housekeeping aaRS's locus tag |
|---|---|---|---|---|
| GCF_017873765.1 | Paenibacillus sediminis | E | J2Z20_RS17205 | J2Z20_RS16290 |
| GCF_000316035.1 | Paenibacillus sp. PAMC 26794 | E | D829_RS0108480 | D829_RS0127060 |
| GCF_016907695.1 | Actinokineospora baliensis | I | JOD54_RS21435 | JOD54_RS29790 |
| GCF_011764545.1 | Phytohabitans flavus | I | Pflav_RS04465 | Pflav_RS27415 |
| GCF_014472405.1 | Micromonospora chalcea | I | H9X95_RS00955 | H9X95_RS23825 |
| GCF_900289095.1 | Pseudomonas syringae | L | DTQ22_RS30290 | DTQ22_RS26320 |
| GCF_003851365.1 | Pseudomonas chlororaphis | L | C4K14_RS13030 | C4K14_RS29135 |
| GCF_007993905.1 | Pseudomonas sp. SJZ077 | L | FBY01_RS12590 | FBY01_RS18180 |
| GCF_002113025.1 | Pseudomonas sp. B28(2017) | L | CAF65_RS05400 | CAF65_RS21805 |
| GCF_902706035.1 | Pantoea sp. 18069 | M | GVL12_RS08265 | GVL12_RS03245 |
| GCF_011683915.1 | Pelagibius litoralis | M | HBA54_RS27400 | HBA54_RS04030 |
| GCF_002366365.1 | Staphylococcus pseudintermedius | R | BSR33_RS06270 | BSR33_RS00740 |
| GCF_015709455.1 | Methylosinus sp. H3A | R | IY145_RS00585 | IY145_RS19355 |
| GCF_021460825.1 | Acinetobacter baumannii | R | LH678_RS08430 | LH678_RS17695 |
| GCF_017166225.1 | Acinetobacter pittii | R | JRG17_RS10170 | JRG17_RS07590 |
| GCF_900452515.1 | Legionella birminghamensis | R | DYH42_RS12890 | DYH42_RS11490 |
| GCF_014872755.1 | Bacillus sp. Bvel1 | R | HAP68_RS17590 | HAP68_RS17540 |
| GCF_022097155.1 | Bacillus subtilis | R | HXV89_RS18970 | HXV89_RS19075 |
| GCF_002998055.1 | Bacillus atrophaeus | R | C6W22_RS18140 | C6W22_RS11480 |
| GCF_016863395.1 | Sphaerisporangium rufum | R | Sru01_RS04260 | Sru01_RS23020 |
| GCF_013393275.1 | Paenarthrobacter nitroguajacolicus | R | DM791_RS11990 | DM791_RS06065 |
| GCF_003399705.1 | Virgibacillus dokdoensis | R | CAI16_RS19440 | CAI16_RS20415 |
| GCF_000214845.1 | Thermus thermophilus | V | THTHE16_RS10075 | THTHE16_RS06030 |

### Supplementary Table 2 | Bacterial strains and vectors used in the study.

| Strain | Res. | Description | Reference or source |
|---|---|---|---|
| *Rhizobium anhuiense* T24 | - | TFX-producing strain, contains the *tfxHABCDEFG* cluster on the chromosome. | USDA collection (USDA2124)[3,4] |
| *R. leguminosarum* 4292 | $Rf^R$ | TFX-sensitive strain, referred to as *Rlp*_4292 in this study. | Common laboratory strain |
| *R. leguminosarum* 4292 *yejA*ΩpVO155 | $Rf^R$, $Km^R$ | Insertion mutant in the *yejA* gene encoding the periplasmic subunit of the YejABEF transporter. | This study |
| *E. coli* Rosetta 2 (DE3) pLysS | $Cm^R$ | Heterologous protein expression. | Novagen |
| *E. coli* BL21 (DE3) | - | Heterologous protein expression. | Novagen |
| *E. coli* DH5α | - | Molecular cloning of all constructs. | Common lab strain |
| **Vector** | **Res.** | **Description** | **Ref. or source** |
| pSRK | $Gm^R$ | Broad-host range vector for inducible (lac promoter) protein expression, oriV (pBBR5 derivative) | Ref.[5] |
| pSRK *tfxD* | $Gm^R$ | *tfxD* gene from the *tfx* BGC of *R. anhuiense* T24 | This study |
| pSRK *tfxE* | $Gm^R$ | *tfxE* gene from the *tfx* BGC of *R. anhuiense* T24 | This study |
| pSRK *tfxG* | $Gm^R$ | *tfxG* gene from the *tfx* BGC of *R. anhuiense* T24 | This study |
| pSRK *tfxH* | $Gm^R$ | *tfxH* gene from the *tfx* BGC of *R. anhuiense* T24 | This study |
| pSRK *Rlp-argRS*$^{HSK}$ | $Gm^R$ | *argRS* gene from *Rlp*_4292 | This study |
| pSRK *Rlp-ileRS*$^{HSK}$ | $Gm^R$ | *ilerS* gene from *Rlp*_4292 | This study |
| pSRK *Lbi-argRS*$^{BGC}$ | $Gm^R$ | *argRS* gene found in the *tfx*-like BGC of *Legionella birminghamensis* | This study |
| pSRK *Lbi-argRS*$^{HSK}$ | $Gm^R$ | *argRS* gene from the genome of *Legionella birminghamensis* outside the BGC | This study |
| pSRK *Eco-argRS*$^{HSK}$ | $Gm^R$ | *argRS* gene from *E. coli* MG1655 | This study |
| pSRK *Sme-yejABEF* | $Gm^R$ | *yejABEF* genes from *S. meliloti* 1021 coding for the components of the YejABEF ABC transporter | Ref.[6] |
| pSRK *Eco-yejABEF* | $Gm^R$ | *yejABEF* genes from *E. coli* MG1655 | Ref.[6] |
| pSRK *Rlp-pepN* | $Gm^R$ | *pepN* gene from *Rlp*_4292 | This study |
| pSRK *Eco-pepN* | $Gm^R$ | *pepN* from *E. coli* MG1655 | This study |
| pET22b(+) | $Ap^R$ | Expression vector for proteins in *E. coli* | Novagen |
| pET22 *tfxH*-CHis6 | $Ap^R$ | Overexpression of TfxH-His6 fusion in *E. coli* | This study |
| pET22 *Rlp-argRS*$^{HSK}$-CHis6 | $Ap^R$ | Overexpression of ArgRS$^{Rle}$-CHis6 fusion in *E. coli* | This study |
| pCA24 NHis6-*Eco-argRS*$^{HSK}$ | $Cm^R$ | Overexpression of NHis6-*Eco*-ArgRS$^{HSK}$ fusion in *E. coli* | ASKA library[7] |
| pET28 NHis6-*Lbi-argRS*$^{HSK}$ | $Km^R$ | Overexpression of NHis6-*Lbi*-ArgRS$^{HSK}$ fusion in *E. coli* | This study |
| pET28 NHis6-*Lbi-argRS*$^{BGC}$ | $Km^R$ | Overexpression of NHis6-*Lbi*-ArgRS$^{BGC}$ fusion in *E. coli* | This study |
| pSRK-pT5-*lac Rlp-argRS*$^{HSK}$ | $Gm^R$ | Overexpression of NStrepII-*Rlp*-ArgRS$^{HSK}$-CHis6 fusion in Rlp_4292 contains the T5-*lac* instead of the original *lac* promoter of the pSRK vector | This study |
| pRK600 | $Cm^R$ | Helper plasmid for conjugal DNA transfer | Ref.[8] |
| pVO155-nptII-gfp | $Ap^R$ | Insertional gene inactivation in rhizobia | Ref.[9] |
| pVO155-nptII-gfp-*Rlp-yejA* | $Ap^R$ | Contains a 563 bp-long internal fragment of the *yejA* gene (RLEG18_RS0119020) from *Rlp*_4292 | This study |
| pVO155-nptII-gfp-*Rlp-pepN* | $Ap^R$ | Contains a 510 bp-long internal fragment of the *pepN* gene (RLEG18_RS0101975) from *Rlp*_4292 | This study |

Ap – ampicillin, Cm – chloramphenicol, Gm – gentamycin, Km – kanamycin, Rf – rifampicin

### Supplementary Table 3 | Nucleotide sequences of primers used in the study.

| Primer name | Primer sequence (5’-3’)* | Purpose |
|---|---|---|
| tfxD_NdeI_F | atatatt**CATATG**agcgacgaaaaccagcatgg | PCR amplification of the *tfxD* gene for cloning into the pSRK vector |
| tfxD_XhoI_R | taaattat**CTCGAG**ttatctgttcggtagtgcatttagggcag | |
| tfxE_NdeI_F | atattaaa**CATATG**cactaccgaacagataaaccg | PCR amplification of the *tfxE* gene for cloning into the pSRK vector |
| tfxE_XhoI_R | taaattat**CTCGAG**tcatgaataactcaccgcctgtatc | |
| tfxE_NdeI_mut_F | cgcaagatcatctgtacgggaaccg | Site-directed mutagenesis to eliminate the internal NdeI restriction site within the *tfxE* gene |
| tfxE_NdeI_mut_R | ggttcccgtacagatgatcttgcgacc | |
| tfxG_NdeI_F | atattat**CATATG**aatgatgagatttgcctgacag | PCR amplification of the *tfxG* gene for cloning into the pSRK vector |
| tfxG_XhoI_R | attattt**CTCGAG**ctatgccagcgctg | |
| tfxH _NdeI_F | atattatt**CATATG**cagcgagaaaacctcgc | PCR amplification of the *tfxH* gene for cloning into the pSRK vector |
| tfxH_XhoI_R | tattatt**CTCGAG**ttaccccatcagctttttttccattttgg | |
| tfxH_nostop_XhoI_R | tattatt**CTCGAG**ccccatcagctttttttccattttgg | PCR amplification of the *tfxH* gene for cloning into the pET22 vector |
| argRS_Rle_NdeI_F | atatattt**CATATG**aaccttttaccgacttcgaag | PCR amplification of the *argRS* gene of *R. leguminosarum* for cloning into the pSRK vector |
| argRS_Rle_XhoI_R | tttatata**CTCGAG**ttatcgcatttcgtccggtgcg | |
| ileRS_Rle_NdeI_F | atattatt**CATATG**accgacacagccgaaaagatc | PCR amplification of the *ileS* gene of *Rlp*_4292 for cloning into the pSRK vector |
| ileRS_Rle_XhoI_R | attaattt**CTCGAG**tcacttgagcgcggcaagctc | |
| argRS_Ec_NdeI_F | atatat**CATATG**aatattcaggctcttctctcagaaaaag | PCR amplification of the *argRS* gene of *E. coli* for cloning into the pSRK vector |
| argRS_Ec_XhoI_R | atatatt**CTCGAG**ttacatacgctctacagtctcaatacccag c | |
| yejA_pVO_SalI_F | ttatat**GTCGAC**caacgggtcaggaccagccc | PCR amplification of the *yejA* gene fragment for cloning into the pVO155-nptII-gfp vector |
| yejA_pVO_XbaI_R | attatta**TCTAGA**tggcttgccgtccggtc | |
| yejA_Rle_check_F | cccggcaaaaaccaaactctctcc | Validation of the *R. leguminosarum yejA* gene inactivation by plasmid insertion |
| yejA_Rle_check_R | ccgggcgagaaggaagcaatc | |
| pepN_pVO_SalI_F | acttcg**GTCGAC**ggcgcggaata | PCR amplification of the *pepN* gene fragment for cloning into the pVO155-nptII-gfp vector |
| pepN_pVO_XbaI_R | tatta**TCTAGA**gatgtcgtcatcgccgcgtaga | |
| pepN_Rle_NdeI_F | ttata**CATATG**cgaacagataccggccaggtcat | PCR amplification of the *pepN* gene of *Rlp*_4292 for cloning into the pSRK vector |
| pepN_Rle_HindIII_R | taatat**AAGCTT**ttaccctttaagcgtgcgctcgacgat | |
| pepN_Eco_NdeI_F | ttatat**CATATG**actcaacagccacaagccaaataccg | PCR amplification of the *pepN* gene of *E. coli* for cloning into the pSRK vector |
| pepN_Eco_HindIII_R | ttatat**AAGCTT**aagccagtgctttagttatcttctcgtac | |
| Str_Rle_argRS_NdeI_F | ttata**CATATG**gcaagctggagtcacccgcagttcgaaaagac tgcGGATCCgaaccttttaccgacttcgaagcc | PCR amplification of the *argRS* gene of *Rlp*_4292 for cloning into the pSRK vector |
| His_Rle_argRS_NheI_R | ttata**GCTAGC**ttagtggtgatggtgatggtgGTCGACtcgca tttcgtccggtgcggcagt | |
| lacI_Mlu_F | gtaaagcggcggtgcacaat | pSRK-T5-lac vector construction |
| T5pro_RNde | ttata**CATATG**taatttctcctctttaatgaattctgtgtgaa | |

*Restriction sites used for cloning are shown with capital bold letters in the sequences of the primers.

### Supplementary Table 4 | Sequences of *lbi-argRS*$^{BGC}$ and *lbi-argRS*$^{HSK}$ genes

| Gene | Nucleotide sequence |
|---|---|
| 6His-*lbi-argRS*$^{BGC}$ | ATGGGCAGCAGC**CATCATCATCATCATCAC**AGCAGCGGCCTGGTGCCGCGCGGCAGCCATATGTCAGAACTATT<br>TGATGACAAAATAAATCAAATTTTTCAAAACGCGTTCGAAATGCTGGGTTATAACACCCAGCATGCAAAGGTCG<br>TGCTGAGCTCTCGCCCTGAACTGGGCCACTTCCAATGTAATGCGGCTATGTCCCTAGCTAAAGAAGTAGGCAAA<br>AACCCGCTGGAAATTGCTAACCACATCGTTGAACTGATCACCAAGAATGAATGGTTTACCAATATCAACATTGC<br>GCGTCCGGGTTTTATCAATTTTAGCGTTAGCAGCGACTACCTGGCGGATTGGTGTAATCAGTTAGTTCTGGATC<br>ATCGTAGCGGTTGCCCGATTGTCGCGAACAAAAAAAAAGTTGTTATCGACTTCGGCGGTCCGAATATTGCCAAG<br>CCGATGCACGTTGGCCATATCCGTAGCACGGTGATCGGCGACTGCCTGCAACGTCTGTATCGTTTCTGCGGTGA<br>CGAGGTTATTTCCGACATCCACCTGGGCGACTGGGGTACACAGATGGGTATGCTGATCGAGGCAATGAAAAAGA<br>AGATGCCGGAAGCGGATTACTTCAATGATGCATTTGAATATGATTACCCGGACAGCTCGCCCGTCACCGTAGAT<br>GAGTTGGCGGCGCTGTACCCGCAGGCTGCGGCGCTGTGCAAGGCGGACGAGATCGAGATGCAGAAAGCGCGTAC<br>CTCTACCCTGGAGCTGCAAAAGAAAAAGCGCGGTTTGGTGGCGCTTTGGACGCATTTCAGCCGCATTAGTATTG<br>AGTCCCTCAAGAAGGACTTCAACGAGCTGGGGGTGACCTTTGACTTGTGGAATGGCGAATCTACGGTGCATGAT<br>TTGGCCGAACGCCTGATCGAGCGCCTGATCGAGAACAAGACTGCGGTCGAATCCAATGGTGCGACCATTATTGA<br>GTTGGAGGAGGATATGCCACCGCTGATACTGAAAAAGAGCGACGGCGCATTCATGTATGGCACGACCGATCTGG<br>CGACCATTGTGGAAAGAGTTAACACCATCAACCCAGATTTGATTTTGTACGTGGTTGATAAACGCCAGAGCCTG<br>CACTTCAAACAGGTTTTTCAGGCTGCGAAAAAGTGCCAGCTGGCGGACTCCGTGATCCTGGAGCACGTTGGTTT<br>TGGTACTGTGAACGGTAGTGACGGCAAACCATTTAAAACCCGTGGTGGCGGTACGATGTCGTTGAAGGATTTGA<br>TCAAAGAAGCAAAGGTGAACGCCGGTAGCCGCTTCAATAACATCGACATCGATGAAAACTTAAAGAACGACATC<br>GTGAACAAGATTGCTCTCAGCACCCTGAAGATTGCGGACCTTTCCAATCCGCGTATTAGCGACTACATTTTCGA<br>CTTGAATGTGGCCTCGCAATTTGAGGGTAAAACTGGTCCGTATCTGCTGTACAGCGCAGTTCGTATCAAAAGCA<br>TGTTGCAAAAAGCCAACAACCTGAACCAGAATGATCTAAAGATCTTACCGGCAGTGAACTTGCTCGAAGAGAAC<br>CTGCAACTGACCTTCACCAAGCTGCCGAAAGCGATTGGCACCGCATACACCAAATTCGAGCCGCACCACTTATG<br>CGATTTTGCATTCGAGTTAGCCTCCGCATATAACCGTTTTTATGCGTCTTGTAATGTTCTGACCGAAGAGAACC<br>TGGTTATCCGTAATAGCAGACTGGCTATGTCTGCCCTGACCCTGAAGCAGATGTTGCTGACTTTCGATATCCTG<br>GGCCTGGAAGTGCCGGATCGTATGTAA |
| 6His-*lbi-argRS*$^{HSK}$ | ATGGGCAGCAGC**CATCATCATCATCATCAC**AGCAGCGGCCTGGTGCCGCGCGGCAGCCATATGAAACAAACGGT<br>GGAGAACCTTCTGGCGGACGCAATTCGCTCGCTCAAAACGAGCGGTGTTATTCCCAGCGACGTGGACGTTGAGA<br>TCAAGGTTGACCGTAGCAAAGATTCGTCGCACGGCGATTTCGCCTCCAACCTCGCACTGATCCTCGCAAAGCCG<br>TGCCGCCAACCCCCGCGCCAGCTCGCCGAACTGCTCATTCAAACGATCGGTCAGCATGAAAGCGTTGAAAAAAT<br>TGAGATTGCCGGTCCGGGCTTTATCAACTTCTTCATGCAGAAAGACTCCCTTTCCCAGGTTGTGGCGACCATCC<br>TTGAGCAACAAGAAAATTACGGTCGCTCCAACCTGGGTAATCAGCAACCCGTTCTGATTGAGTTCGTTTCCGCC<br>AATCCCACCGGCCCCCTGCATGTGGGTCACGGTCGCGGTGCGGCGTTTGGTGCAACCCTGGGTAATCTGCTCTC<br>CGCCGCGGGCTACAAGGTGAGCCGCGAGTATTATGTGAACGACGCAGGTCGTCAAATGAATATCCTTGCCGTGT<br>CCGTGTGGCTTCGGTATCTTACCCTTACGGGTGAATCGATTATTTTCCCCGCCAATGGCTATCGCGGTGATTAT<br>GTGAACGATATTGCACGCATGGTCTTTGAGAGCCACGGTGAAAAATATAAACAACCCTGGGGCCTTATCAAGGA<br>GTCCCTCCCGGCAGACGAACCGGAGGGTGGCGACAAAGAAGTCTATATTGATGCGGTGATCGAACGCTGCAAAC<br>AGCTTCTGGGTCTTGACGGTTTTAAATACTTTCACAAGGAGGCGCTGGACTCCGTCCTCGATGACATTAAAGAC<br>GACCTTGAGGATTTTGGCGTTCACTTCGATTGCTGGTTCTCCGAGCAACACCTGCTGGAGGACGGTTCGATTGA<br>GAAAGGCATTCAGGCCCTCAAAGACGCGAACCACACCTACACCAAGGAGGGTGCACTCTGGTTTAAGGCAACCG<br>ATTTTGGCGATGAAAAAGATCGCGTTCTCATTCGCGCCAATGGCCAGACGACCTACTTCGCGAGCGATGTTGCA<br>TATCATTGGAATAAATACGATCGCGGTTTCGCCCGCGTCATTGATATTTTTGGCGCGGACCACCACGGCTACAT<br>CAGCCGGGTGCGGGCGGCGGTCAATGCACTCGGCCACGACGATAATGCACTGAACGTTCTTCTTGTGCAATTCG<br>CGATCCTGTATCGGGGTGGCCATCGGGTGCAGATGTCGACGCGCAGCGGTAGCTTCGTCACCCTTCGCGAGCTC<br>CGGGAAGAAGTCGGTAAGGACGCAGCGCGCTTCTTTTATGTCATGCGCAAGCCGGAGCAACACATGGATTTTGA<br>CCTTGACCTCGCGAAGAGCCAAAGCTCCGACAATCCCGTCTACTACATCCAGTACGCGCACGCCCGTATTTCGT<br>CCGTCCTCCGCCAACTCAAAGACCGGGGTCTTTCGTGGTCGGAGGAGATCGGTCTGGCCAACATTGCAAAACTG<br>GTTGAACCGCATGAGATCTCCCTTATGACGCTTATTTCCCGTTATCCGGAAGTTATCGTTTCCGCGGCCAACAC<br>GTGTGAGCCCCACCAAATCGCCTACTACCTTCGTGAACTGGCGAATGGTCTCCATTCGTATTATAACGCCGTCA<br>CGCTCCTTTGCGAACAGGACACGCTCCGCAGCGCCCGGCTGAGCCTTCTTACCGCAGTCCGGCAAGTGCTGCGT<br>AATGGCCTCCAACTTCTCGGCGTCAGCTGTCCCGAGTCCATGTAA |
